# Open Soil Spectral Library (OSSL): Building reproducible soil calibration models through open development and community engagement

**DOI:** 10.1101/2023.12.16.572011

**Authors:** José L. Safanelli, Tomislav Hengl, Leandro Parente, Robert Minarik, Dellena E. Bloom, Katherine Todd-Brown, Asa Gholizadeh, Wanderson de Sousa Mendes, Jonathan Sanderman

## Abstract

Soil spectroscopy is a widely used method for estimating soil properties that are important to environmental and agricultural monitoring. However, a bottleneck to its more widespread adoption is the need for establishing large reference datasets for training machine learning (ML) models, which are called soil spectral libraries (SSLs). Similarly, the prediction capacity of new samples is also subject to the number and diversity of soil types and conditions represented in the SSLs. To help bridge this gap and enable hundreds of stakeholders to collect more affordable soil data by leveraging a centralized open resource, the Soil Spectroscopy for Global Good has created the Open Soil Spectral Library (OSSL). In this paper, we describe the procedures for collecting and harmonizing several SSLs that are incorporated into the OSSL, followed by exploratory analysis and predictive modeling. The results of 10-fold cross-validation with refitting show that, in general, mid-infrared (MIR)-based models are significantly more accurate than visible and near-infrared (VisNIR) or near-infrared (NIR) models. From independent model evaluation, we found that Cubist comes out as the best-performing ML algorithm for the calibration and delivery of reliable outputs (prediction uncertainty and representation flag). Although many soil properties are well predicted, total sulfur, extractable sodium, and electrical conductivity performed poorly in all spectral regions, with some other extractable nutrients and physical soil properties also performing poorly in one or two spectral regions (VisNIR or Neospectra NIR). Hence, the use of predictive models based solely on spectral variations has limitations. This study also presents and discusses several other open resources that were developed from the OSSL, aspects of opening data, current limitations, and future development. With this genuinely open science project, we hope that OSSL becomes the driver of the soil spectroscopy community to accelerate the pace of scientific discovery and innovation.

## Introduction

The need for high-quality soil data has grown exponentially in recent years. Monitoring soils for sustainable development is a high priority at this moment as a number of large existential environmental challenges have been linked to the role that soils play [1, 2]. Healthy soils are essential for the growth of crops, filtration of water, functioning of ecosystems, and storage of vast amounts of carbon, with this latter point being not only crucial for climate change mitigation potential but also for sustainable land stewardship, ultimately contributing to improved agricultural and environmental outcomes [3–5]. In turn, soil scientists have been struggling to meet this demand because measuring soil properties still relies largely on physical sampling and resource-intensive benchtop (wet-chemistry) analytical methods [6]. Similarly, many investigations and modeling of the global challenges of soil security still rely on limited soil expertise or legacy data, which is not compatible with the urgent needs [7, 8]. Tackling the challenge of meeting society’s demands for soil information has been proposed with strong community-based research efforts and the incorporation of novel (or not so) technologies, like dry-chemistry soil spectroscopy [6, 7].

Diffuse reflectance spectroscopy, the measurement of light absorption in the visible and near-infrared (VisNIR; 350–2500 nm) or mid-infrared (MIR; 2500–25000 nm [4000–400 cm^*−*1^]) regions of the electromagnetic spectrum, has emerged as an important, rapid and low-cost complement to traditional wet chemical analyzes [9–11]. The proportion of incident radiation reflected by the soil is detected through VisNIR-MIR reflectance spectroscopy and can be used to estimate numerous soil properties using statistical and machine learning methods [12–14]. However, a bottleneck to the more widespread adoption of soil spectroscopy is the need for establishing large reference training data sets, which are called soil spectral libraries (SSLs) [11]. In fact, for dealing with complex relationships between soil components and the spectra, irrespective of the multivariate statistical or machine learning method (e.g., local or global modeling), the prediction capacity for new soil samples is still subject to the number (quantity) and diversity (quality) of soil types and conditions that are represented in the feature space (spectra), a limitation that is not exclusive to soil spectroscopy but rather common to any machine learning problem [15].

In recent years, there were some attempts to build a global soil spectral library based on collaborative contributions that could serve the soil science community, e.g. by Viscarra-Rossel et al. [13] and Demattê et al. [16]. More recently, the Global Soil Partnership of the Food and Agriculture Origination (FAO) of the United Nations and its Global Soil Laboratory Network initiative on soil spectroscopy (GLOSOLAN-Spec) started engaging the international soil science community to connect conventional soil testing laboratories to those having or interested in having spectroscopy capacity. An unprecedented effort to establish standards and protocols for soil spectroscopy has emerged as part of the Institute of Electrical and Electronics Engineers (IEEE) Standards Association (AS) P4005 working group. Nonetheless, a step towards community integration that genuinely observes open-data principles was lacking at the moment.

To bridge this data gap and enable hundreds of stakeholders interested in soil information generation by leveraging a centralized open resource, the Soil Spectroscopy for Global Good (SS4GG) initiate has created an open-source and open-data project called the *“Open Soil Spectral Library”* (OSSL). The SS4GG was designed to accelerate the pace of scientific discovery in soil spectroscopy by facilitating and supporting a collaborative network of researchers. With the OSSL, the SS4GG team seeks to develop advanced yet intuitive, open-source, web-hosted resources to predict various soil properties from spectra collected on any spectrometer anywhere in the world. This initiative brings together soil scientists, spectroscopists, pedometricians, data scientists, and others to overcome some of the current bottlenecks preventing wider and more efficient use of soil spectroscopy. A series of events and activities have been initiated to address topics including lab-to-lab interoperability [17], data harmonization, model development, community engagement & outreach, and the use of spectroscopy to inform environmental modeling (https://soilspectroscopy.org/).

This paper presents a detailed and technical description of the OSSL and the resources derived that are currently available to the soil spectroscopy community. The OSSL is based primarily on the work and ideas of Viscarra-Rossel et al. [13] and Shepherd et al. [6]; it can be considered a *scale-up* of the first global soil calibration library produced by ICRAF-ISRIC [18]. Here, we describe the methods for collecting and harmonizing several SSLs that are incorporated into the OSSL database. We then provide an exploratory data analysis of the combined data and report the accuracy of the fitted prediction models built on top of the OSSL database. All processing steps and the modeling inputs and outputs are fully documented on Github repositories (https://github.com/soilspectroscopy) and the OSSL technical manual (https://soilspectroscopy.github.io/ossl-manual/). In this paper, we provide only key results from the accuracy assessment and discuss possible limitations and future development directions. The OSSL is an ongoing project, being periodically updated with new data and model re-calibration.

## Materials and methods

### Preparing the OSSL database

The OSSL is made up of several SSLs that present a data-sharing policy. The latest version (v1.2) includes the United States Department of Agriculture (USDA) Natural Resources Conservation Service (NRCS) National Cooperative Soil Survey (NCSS) - Kellogg Soil Survey Laboratory (KSSL) MIR [19–21] and VisNIR libraries, labeled `KSSL`, with the VisNIR dataset being sourced from the Rapid Carbon Assessment Project (RaCA) project from USDA-NRCS [22, 23]; World Agroforestry Centre (ICRAF) MIR and VisNIR libraries, developed in cooperation with the International Soil Reference and Information Centre (ISRIC) [24], labeled `ICRAF-ISRIC`; two MIR libraries of the Africa Soil Information Service (AfSIS) [25, 26], labeled as `AFSIS1` and `AFSIS2`; the VisNIR library from the European Data Centre (ESDAC) — Joint Research Centre (JRC) as part of the Land Use/Cover Area frame statistical Survey (LUCAS soil of 2009, 2012, and 2015) [27], labeled `LUCAS`; the Central African MIR SSL developed by ETH Zurich in the Congo Basin [28], labeled `CAF`; a MIR library of New Zealand forest soils from researchers of Scion Research, NZ [29], labeled `Garrett`; a MIR library of boreal soils from researchers of the University of Zurich [30], labeled `Schiedung`; and a subset of the Serbian SSL from the University of Novi Sad [31], labeled `Serbia`.

The KSSL MIR dataset represents a snapshot from July 2022, as this large SSL keeps growing in size as more samples are scanned. Aliquots of around 600 samples from the LUCAS SSL were shipped to Woodwell Climate Research Center and scanned using an MIR device (Bruker Vertex 70 with a Pike Autodiff accessory). In terms of data licensing, `KSSL`, `ICRAF-ISRIC`, `AFSIS2`, `CAF`, `Schiedung`, and `Garrett` present a Creative Commons (CC) license, while LUCAS has a JRC license agreement with *“(*…*) graphical representation of individual units on a map is permitted as long as the geographical locations of the soil samples are not detectable”* (i.e., downgraded to about 1 km of precision), `AFSIS1` has an Open Data License (ODbL), and `Serbia` was published in a paper and shared internally with the OSSL team.

In addition to the full VisNIR and MIR spectral ranges, we imported into the OSSL a library limited to a specific NIR range (1350–2550 nm) built using the NeoSpectra Handheld NIR Analyzer developed by Si-Ware. This library includes 2,106 distinct mineral soil samples scanned on 9 portable low-cost NIR spectrometers identified by serial number. 2,016 of these soil samples were selected to represent the diversity of mineral soils found in the United States, and 90 samples were selected across Ghana, Kenya, and Nigeria. The 2,016 soil samples were obtained from USDA NRCS NSSC-KSSL Soil Archives and scanned by the Woodwell Climate Research Center and the University of Nebraska — Lincoln, which were made available with a CC-BY 4.0 license on Zenodo [32]. The number of unique samples correctly imported into the OSSL per original source and type of spectra is described in Table 1. A diagram of the general OSSL workflow is provided in Fig. 1.

**Table 1.** Number of unique samples correctly imported into the OSSL per original source and spectra type.

| SSL | VisNIR | MIR | NIR |
| --- | --- | --- | --- |
| ICRAF-ISRIC | 4073 | 4071 |  |
| KSSL | 19807 | 76813 |  |
| LUCAS | 40764 | 589 |  |
| AFSIS1 |  | 1904 |  |
| AFSIS2 |  | 151 |  |
| CAF |  | 1578 |  |
| Garrett |  | 184 |  |
| Schiedung |  | 259 |  |
| Serbia |  | 135 |  |
| Neospectra |  |  | 2106 |

SSL: Soil spectral library; VisNIR: Visible and near-infrared region (350–2500 nm); MIR: Mid-infrared region (600–4000 cm<sup>-1</sup>); NIR–Neospectra: Near-infrared region of the Neospectra device (1350–2550 nm); ICRAF-ISRIC: library from World Agroforestry Centre (ICRAF) developed in cooperation with the International Soil Reference and Information Centre (ISRIC); KSSL: library from the United States Department of Agriculture (USDA) — Natural Resources Conservation Service (NRCS) — National Cooperative Soil Survey (NCSS) — Kellogg Soil Survey Laboratory (KSSL); LUCAS: library from the European Data Centre (ESDAC) as part of the Land Use/Cover Area frame statistical Survey (LUCAS soil of 2009, 2012, and 2015), with MIR spectra scanned at Woodwell Climate Research Center; AFSIS1 and AFSIS2: library of the Africa Soil Information Service (AfSIS); CAF: Central African library developed by ETH Zurich in the Congo Basin; GARRET: library from researchers of Scion Research, NZ; SCHIEDUNG: library produced by the researchers of the University of Zürich; SERBIA: library produced by the University of Novi Sad, Serbia; Neospectra: library based on Neospectra Handheld NIR Analyzer developed by Si-Ware.

**Fig 1.**
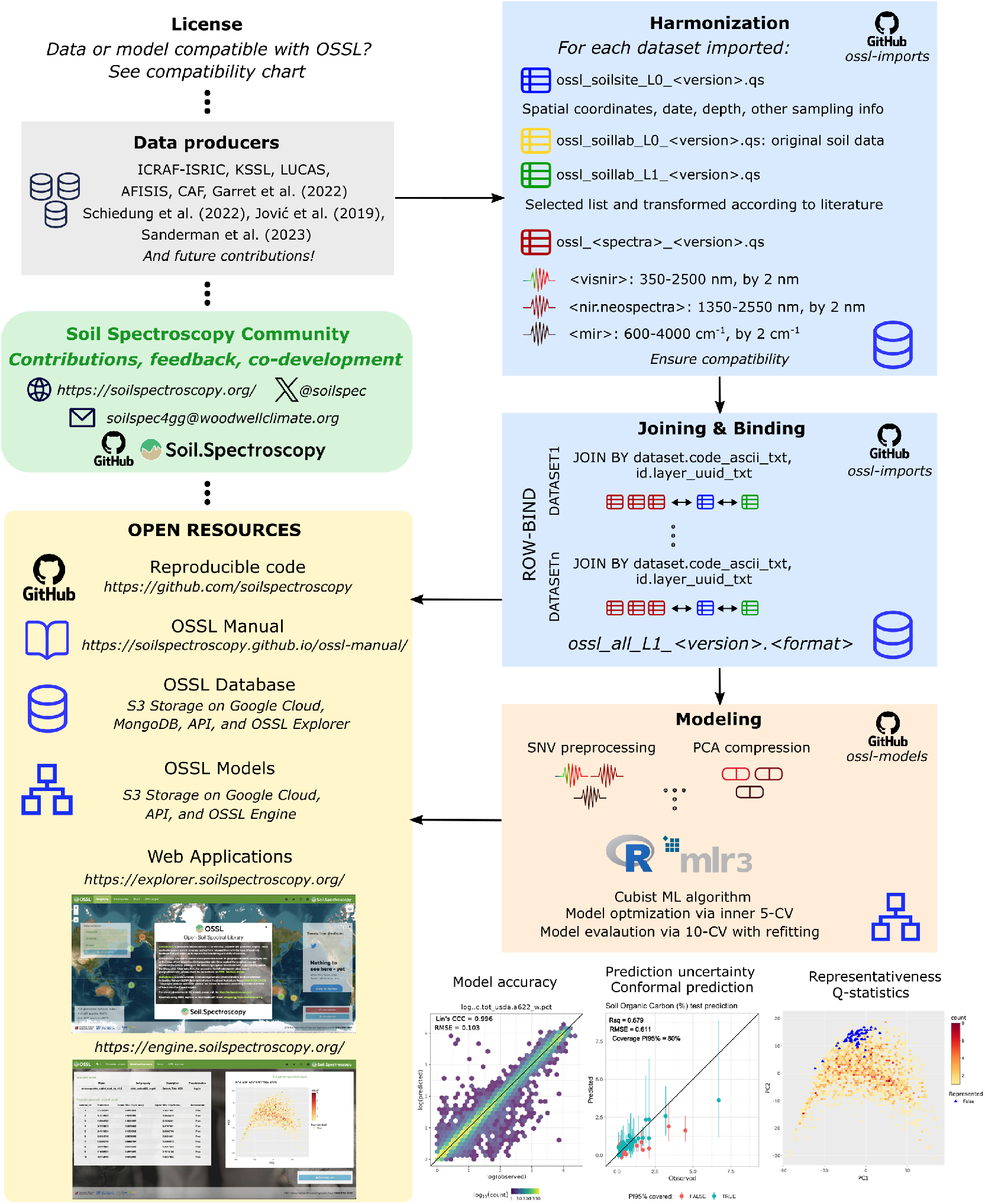
The general processing diagram used to create OSSL. Compilation and harmonization of the Open Soil Spectral Library (OSSL) from original data producers, and modeling framework that includes model calibration, evaluation, additional outputs, and open resources.

### Harmonizing the SSLs

Different SSLs may cause prediction bias in external samples [33] for a number of reasons including different analytical reference methods and standard operating procedures (SOPs) [34], or differences in scanning conditions affected by the instruments themselves and the SOPs used for scanning [35]. Contrasting methods used to determine (i.e., wet chemistry) a given soil property have been a subject of ongoing discussion. Global efforts such as the Global Soil Laboratory Network (GLOSOLAN) [36] grapple with this same challenge in their soil databases, but there is still no clear consensus on how to harmonize those different analytical methods.

To maximize transparency, we have decided to produce two different levels for the analytical reference methods in the OSSL database. Level 0 takes into account the original methods employed in each data set, but initially tries to fit them to two reference guides: i) the KSSL Guidance — Laboratory Methods and Manuals [37]; ii) and any method cataloged by ISO standard [38, 39]. If an analytical method does not fall into any previous methods and has a significant number of samples, then we create a new variable. The final harmonization takes place in OSSL level 1, where those common properties sharing different methods are converted to a target using some publicly available transformation rule, or in the worst scenario, they are naively bound or kept separated to maintain their specificity. The processing code is documented in the *ossl-imports* GitHub repository (https://github.com/soilspectroscopy/ossl-imports) allowing for community-based co-development.

For importing SSLs into the OSSL, we define a schema where soil site information, soil laboratory data (i.e., wet-chemistry), and spectroscopic data are prepared as separate tables that share a common column ID for proper joining. For column names in soil laboratory data, we concatenate three different pieces of information separated by an underscore: abbreviation of the soil property name, analytical method code, and unit abbreviation. For example, plant available calcium (mg kg^*−*1^) using Mehlich 3 as the extractor, following the KSSL procedure, was named `ca.ext_usda.a1059_mg.kg.` Organic carbon (%) determined following ISO 10694 was formatted as `oc_iso.10694_w.pct`. This string pattern is followed in other files and column names with some minor adaptations where required. Reference tables with column names, abbreviations, units, data type, descriptions, and examples are provided in the `ossl-imports` GitHub repository (`ossl_level0_names_soillab.csv, ossl_level0_names_soilsite.csv, ossl_level0_names_visnir.csv, ossl_level0_names_mir.csv`). A reference table containing a copy of the KSSL Guidance procedures is also provided in that repository (`kssl_procedures.csv`). Lastly, the harmonization rules used to translate or combine the soil properties from level 0 to level 1 are also available there (see table: `ossl_level0_to_level1_soillab_harmonization.csv`). Both levels are available in the OSSL database; however, only a selection of harmonized soil properties (level 1) containing at least 500 samples was used to calibrate the prediction models (Table 2).

**Table 2.** Number of samples for the selected and harmonized soil properties of the Open Soil Spectral Library, categorized per spectra and subset type.

| Soil property code | Description | KSSL |  | OSSL |  |  |
| --- | --- | --- | --- | --- | --- | --- |
|  |  | VisNIR | MIR | VisNIR | NIR | MIR |
| acidity_usda.a795_cmolc.kg | Acidity, BaCl<sub>2</sub>-TEA, pH 8.2 |  | 28550 | 1511 | 1576 | 30061 |
| aggstb_usda.a1_w.pct | Aggregate Stability |  | 3218 |  | 197 | 3218 |
| al.dith_usda.a65_w.pct | Aluminum, Crystalline |  | 31135 |  | 1773 | 31135 |
| al.ext_usda.a1056_mg.kg | Aluminum, Extractable, Mehlich-3 |  | 1534 |  | 76 | 3773 |
| al.ext_usda.a69_cmolc.kg | Aluminum, Extractable, KCl |  | 14169 | 1541 | 716 | 15710 |
| al.ox_usda.a59_w.pct | Aluminum, Amorphous |  | 28260 |  | 1354 | 28260 |
| awc.33.1500kPa_usda.c80_w.frac | Available Water Content, 33-1500 kPa |  | 16175 |  | 931 | 16175 |
| b.ext_mel3_mg.kg | Boron, Extractable, Mehlich-3 |  |  |  |  | 2093 |
| bd_usda.a4_g.cm3 | Bulk Density, <2mm fraction | 19752 | 40136 | 20751 | 1085 | 41484 |
| c.tot_usda.a622_w.pct | Carbon, Total NCS | 19807 | 76595 | 19807 | 1976 | 80698 |
| ca.ext_usda.a1059_mg.kg | Calcium, Extractable, Mehlich-3 |  | 1534 |  | 76 | 3773 |
| ca.ext_usda.a722_cmolc.kg | Calcium, Extractable, NH<sub>4</sub>OAc | 51 | 53232 | 3723 | 1976 | 57085 |
| caco3_usda.a54_w.pct | Carbonate, <2mm Fraction | 9213 | 27476 | 51094 | 665 | 29225 |
| cec_usda.a723_cmolc.kg | CEC, pH 7.0, NH<sub>4</sub>OAc | 51 | 53227 | 22687 | 1976 | 57651 |
| cf_usda.c236_w.pct | Coarse Fragments, Greater 2mm |  | 55881 | 23237 | 1969 | 56470 |
| clay.tot_usda.a334_w.pct | Clay | 166 | 50007 | 27145 | 1976 | 57268 |
| cu.ext_usda.a1063_mg.kg | Copper, Extractable, Mehlich-3 |  | 1534 |  | 76 | 3772 |
| ec_usda.a364_ds.m | Electrical Conductivity, (w/w) | 51 | 31886 | 21833 | 942 | 34040 |
| fe.dith_usda.a66_w.pct | Iron, Crystalline |  | 31138 |  | 1773 | 31138 |
| fe.ext_usda.a1064_mg.kg | Iron, Extractable, Mehlich-3 |  | 1534 |  | 76 | 3773 |
| fe.ox_usda.a60_w.pct | Iron, Amorphous |  | 28259 |  | 1354 | 28259 |
| k.ext_usda.a1065_mg.kg | Potassium, Extractable, Mehlich-3 |  | 1534 |  | 76 | 3772 |
| k.ext_usda.a725_cmolc.kg | Potassium, Extractable, NH<sub>4</sub>OAc | 51 | 53230 | 44489 | 1976 | 57674 |
| mg.ext_usda.a1066_mg.kg | Magnesium, Extractable, Mehlich-3 |  | 1534 |  | 76 | 3773 |
| mg.ext_usda.a724_cmolc.kg | Magnesium, Extractable, NH<sub>4</sub>OAc | 51 | 53232 | 3731 | 1976 | 57093 |
| mn.ext_usda.a1067_mg.kg | Manganese, Extractable, Mehlich-3 |  | 1534 |  | 76 | 3772 |
| mn.ext_usda.a70_mg.kg | Manganese, Extractable, KCl |  | 14166 |  | 716 | 14166 |
| n.tot_usda.a623_w.pct | Nitrogen, Total NCS | 19806 | 76594 | 60570 | 1976 | 81282 |
| na.ext_usda.a1068_mg.kg | Sodium, Extractable, Mehlich-3 |  | 1534 |  | 76 | 3616 |
| na.ext_usda.a726_cmolc.kg | Sodium, Extractable, NH<sub>4</sub>OAc | 51 | 53230 | 3715 | 1976 | 57075 |
| oc_usda.c729_w.pct | Organic Carbon, without CaCO<sub>3</sub> | 19747 | 75929 | 64211 | 1974 | 82573 |
| p.ext_usda.a1070_mg.kg | Phosphorus, Extractable, Mehlich-3 |  | 25602 |  | 754 | 27690 |
| p.ext_usda.a270_mg.kg | Phosphorus, Extractable, Bray1 |  | 7146 |  | 266 | 7282 |
| p.ext_usda.a274_mg.kg | Phosphorus, Extractable, Olsen |  | 15359 | 40764 | 281 | 16084 |
| ph.cacl2_usda.a481_index | pH, 1:2 Soil-CaCl<sub>2</sub> Suspension | 50 | 52946 | 40848 | 1976 | 53819 |
| ph.h2o_usda.a268_index | pH, 1:1 Soil-Water Suspension | 50 | 52867 | 44590 | 1976 | 59997 |
| s.tot_usda.a624_w.pct | Sulfur, Total NCS | 19807 | 76592 | 19807 | 1976 | 76592 |
| sand.tot_usda.c60_w.pct | Sand, Total | 166 | 50012 | 27086 | 1976 | 57162 |
| silt.tot_usda.c62_w.pct | Silt, Total | 166 | 50013 | 27139 | 1976 | 57215 |
| wr.1500kPa_usda.a417_w.pct | Water Retention, 15 Bar (1500 kPa) |  | 40284 | 971 | 1951 | 41345 |
| wr.33kPa_usda.a415_w.pct | Water Retention, 1/3 Bar (33 kPa) |  | 18536 | 923 | 1032 | 19459 |
| zn.ext_usda.a1073_mg.kg | Zinc, Extractable, Mehlich-3 |  | 1534 |  | 76 | 3760 |

VisNIR: Visible and near-infrared region (350–2500 nm); MIR: Mid-infrared region (650–4000 cm<sup>-1</sup>); NIR: Near-infrared region from Neospectra (1350–2550 nm).

For the spectral data (Fig. 2), VisNIR spectra were formatted to reflectance factor per wavelength. The spectra imported into the OSSL were trimmed between 350 and 2500 nm, with an interval of 2 nm. MIR spectra were formatted to pseudo-absorbance units per wavenumber. The spectral range imported into OSSL falls between 600 and 4000 cm^*−*1^, with an interval of 2 cm^*−*1^. All data sets followed these specifications, and for some cases, we converted the reflectance (*R*) values into absorbance units (*A*) with *A* = *log*_10_(1*/R*). For measurements having a shorter or larger spacing interval, we used splines for re-sampling the spectral curves into the defined specifications [40].

**Fig 2.**
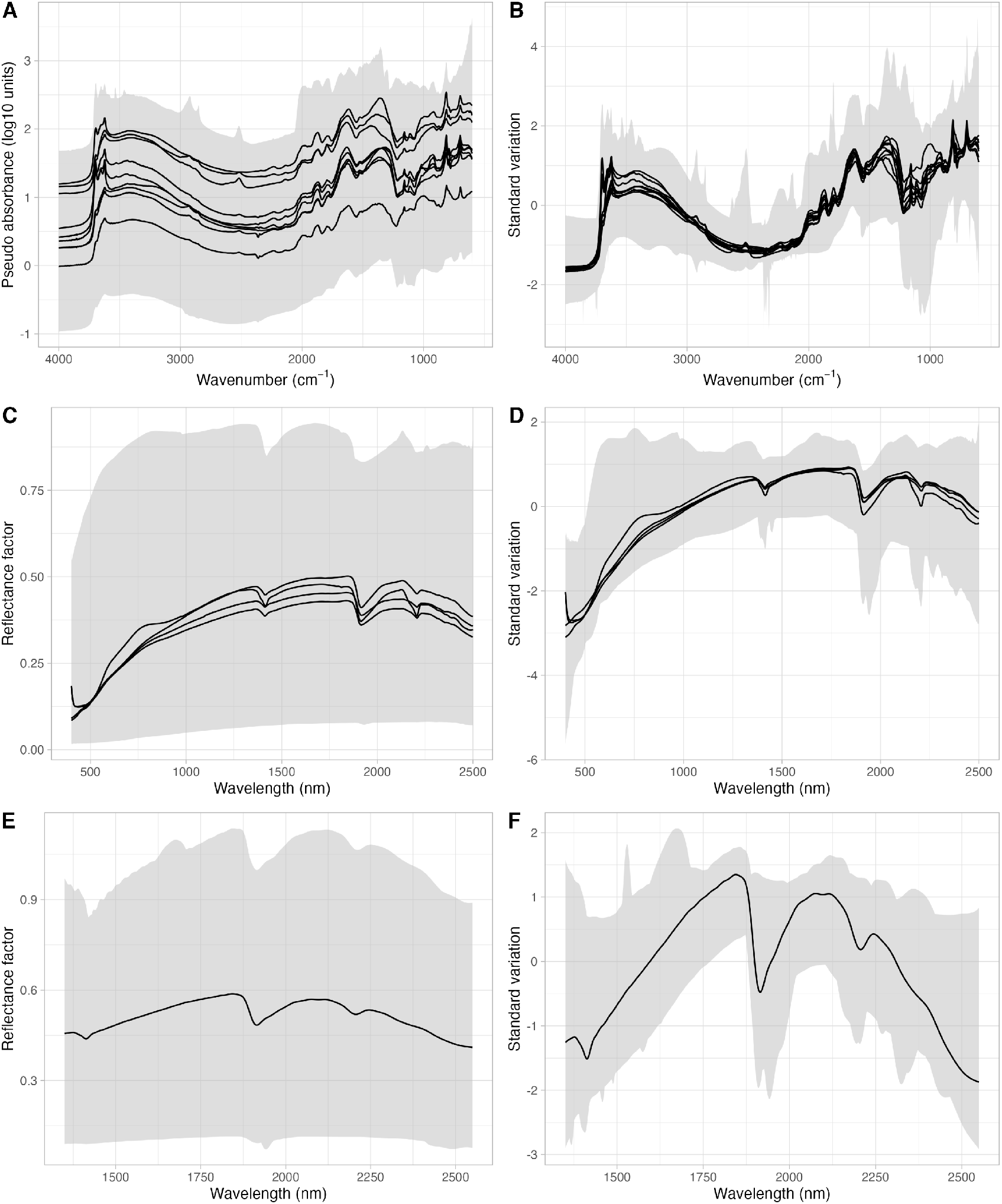
Spectral variation of the mid-infrared (650–4000 cm^*−*^1, panels A and B), visible and near-infrared (350–2500 nm, panels C and D), and near-infrared from Neospectra (1350–2550 nm, panels E and F) imported into the Open Soil Spectral Library (OSSL). The shaded area depicts the maximum and minimum values of the spectral region. The black lines represent the mean of each original dataset imported into the OSSL. Left panels (A, C, and E) are the original spectra, while the right panels (B, D, and F) represent the spectra after preprocessing with Standard Normal Variate (SNV).

### Modeling framework

We have used the MLR3 framework [41] to fit machine learning models. MLR3 is a package universe from R statistical programming language where the user can seamlessly define a machine learning pipeline for resampling, algorithm and model selection, optimization, evaluation, and other steps. In turn, for machine learning, Cubist (*cubist* acronym) [42, 43] was adopted after several studies supporting its use in soil spectroscopy [17, 44, 45] and after the positive findings from the internal benchmarks, which are presented later in this paper. Cubist is an ML algorithm that takes advantage of a decision-tree splitting method but fits linear regression models at each terminal leaf. It also uses a boosting mechanism (sequential trees adjusted by weight) that allows for the growth of a forest by tuning the number of committees. We have not used the correction of final predictions by the influence of nearest neighbors due to the lack of this feature in the MLR3 framework at the time of building OSSL.

The spectral dissimilarity across the original datasets that comprise the OSSL was handled by mathematical pre-processing with standard normal variate (SNV). A ring trial experiment carried out as part of the SS4GG initiative [17] found that SNV is capable of reducing some of the variability caused by the contrasting instruments and SOPs (Fig. 2), in addition to minimizing particle size, light scattering, and diffuse reflectance multicollinearity issues [46]. In turn, spectral standardization may be necessary when spectral responses are very distinct, but this was not applied in this study due to the absence of shared standard samples across laboratories from the imported libraries.

In terms of model types, we have fitted five different versions depending on the availability of samples in the database, relying solely on spectral variation to maximize applicability to different users (`na` acronym), i.e., site information or external environmental layers were not used as ancillary predictors, as they would require spatial and date information in the inputs. The model types were also composed of two different subsets, that is, using KSSL alone (`kssl` acronym), to eliminate potential errors due to analytical data being collected in different ways, or the full OSSL database (`ossl` acronym), in combination with three separate spectral types: VisNIR (`visnir` acronym), MIR (`mir` acronym), and NIR from the Neospectra instrument (`nir.neospectra` acronym). Only the `nir.neospectra` have a single model (`ossl`) as the differentiation would not be advantageous due to the limited sample size (Table 3).

**Table 3.** Modeling combinations fitted with the OSSL database.

| Spectra type | Model subset | Ancillary | Model name |
| --- | --- | --- | --- |
| MIR | kssl | na | mir_cubist_kssl_na.v1.2 |
| MIR | ossl | na | mir_cubist_ossl_na.v1.2 |
| VisNIR | kssl | na | visnir_cubist_kssl_na.v1.2 |
| VisNIR | ossl | na | visnir_cubist_ossl_na.v1.2 |
| NIR–Neospectra | ossl | na | nir.neospectra_cubist_ossl_na.v1.2 |

MIR: Mid-infrared region (650–4000 cm<sup>-1</sup>); VisNIR: Visible and near-infrared region (350–2500 nm); NIR–Neospectra: near-infrared region from Neospectra (1350–2500 nm); kssl: model built solely with the Kellogg Soil Survey Laboratory (KSSL) dataset; ossl: model built with the full Open Soil Spectral Library (OSSL) database; na: site or environmental layers not added as ancillary predictors; cubist: machine learning algorithm.

Hyperparameters were optimized through an internal (inner) re-sampling process using 5-fold cross-validation and smaller subsets (^*∼*^2000 samples) to speed up computing time [47]. This task was carried out with a grid search of the hyperparameter space testing up to five (5) Cubist configurations (*commitees* = [1, 5, 10, 15, 20]) to find the lowest root mean squared error (RMSE, Eq. 4). The final model with the best optimal hyperparameters was fitted at the end with the full training data.

As predictors, we have used the first 120 principal components (PCs) of each spectral range and subset to maximize the balance i.e. trade-off between the spectral representation and compression magnitude employed by principal component analysis (PCA) [48, 49]. Each combination of models in Table 3) had a separate PCA model (Eq. 1) fitted to compress the spectra of the training set and new samples. Additionally, as the first 120 components represent around 99% of the original variability, the remaining 1% was used to verify the extrapolation capacity of the calibration (see below), as this minor amount might contain specific absorption characteristics of unique soil samples that can make the calibration set unrepresentative and affect the predictive capacity.

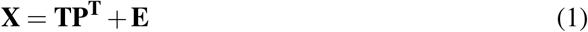

where **X** is the original spectra that are decomposed into the PCA space with new coordinates denominated scores (**T**), estimated by the linear combination of loadings (**P**).

Considering that only the first 120 PCs are used to decompose **X**, the remaining higher-order components are denominated error (**E**), which is calculated by the difference between **X** and 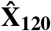. This reasoning was employed in the OSSL models to identify potential underrepresented or outlier samples relative to the calibration set **T**_120_. In this task, each new spectrum (*x*_*i*_) to be predicted is back-transformed using only the first 120 PCs 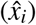. With the back-transformed version 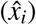, the residual of each sample (*e*_*i*_ = *x*_*i*_ − 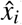) is used to calculate the *Q* statistic (Eq. 2). The *Q*_*i*_ statistic is compared against a critical value (upper confidence limit, *UCL*_*Q*_) estimated across the calibration set with a 99% confidence level (Eq. 3) for identifying potential underrepresented samples when *Q*_*i*_ *> UCL*_*Q*_ [48, 50, 51].

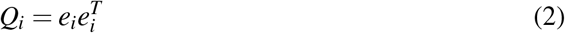

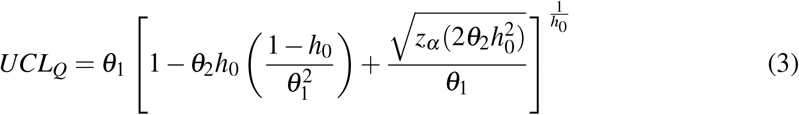

where *θ*_1_ is the trace (sum of the values on the main diagonal) of the covariance of **E**, defined as *Cov*(**E**), *θ*_2_ and *θ*_3_ are the trace of *Cov*(**E**)^2^ and *Cov*(**E**)^3^, respectively, 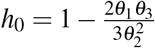, and *z*_*α*_ is the standardized normal variable with (1 *−α*) confidence level (set 99%).

Except for clay, silt, sand, pH H_2_O, and pH CaCl_2_, all other soil properties were natural-log transformed (with offset = 1, `log1p()` R function) to improve the prediction performance of soil properties with highly skewed distribution [44]. They were back-transformed (`expm1()` R function) only at the end after running all the modeling steps, including performance estimation and definition of the uncertainty intervals.

The evaluation of the models was performed with a 10–fold external cross-validation (outer) of the tuned models using RMSE (Eq.4), mean error (bias, Eq.5), R-squared (R^2^, Eq.6), Lin’s concordance correlation coefficient (CCC, Eq.7), and the ratio of performance to the interquartile range (RPIQ, Eq.8) [52]:

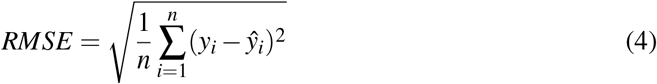

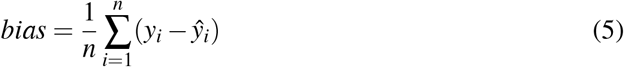

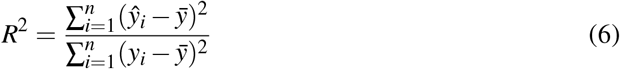

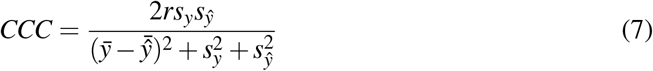

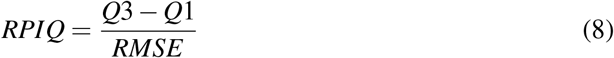

where *y*_*i*_ is the observed value, *ŷ*_*i*_ is the predicted value, 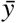is the mean value, *n* is the total number of samples, *r* is the Pearson correlation between *y* and 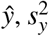 is the variance of observed values, 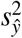 is the variance of predicted values, 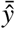 is the mean of predicted values, and *Q*3 *−Q*1 is the interquartile distance of observed values.

Cross-validated predictions were also used to estimate the unbiased absolute error (*δ*_*i*_ = |*y*_*i*_ *− ŷ*_*i*_|), which is further used to calibrate the uncertainty of the estimation models via conformal prediction [53]. The conformal prediction is based on past experience, that is the error model (*δ* = *f* (**T**_120_)) is calibrated using the same fine-tuned structure of the respective response model (*y* = *f* (**T**_120_)). Non-conformity scores (*α*) were estimated from the predicted calibration error with a confidence level of 68% (*ε*) to approximate one standard deviation (Eq. 9). Non-conformity scores from the calibration set are sorted in increasing order to identify the percentile corresponding to the predefined confidence level (*α*_*ε*_). Finally, for new samples, the prediction interval (*PI*) can be derived from the predicted response (*ŷ*) and the predicted error 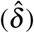 corrected by the non-conformity score of the calibration set (Eq.10) [54]. The main advantage of conformal prediction is the coverage of the intervals; that is, a confidence of 80% means that the predicted range may contain the observed value for around or more than 80% of the cases [53]. Another big advantage of conformal prediction methods is that we get rid of re-sampling mechanisms such as bootstrapping or Monte Carlo simulation, which require a huge computation cost for large sample sizes and may not have good coverage of the true estimates. The non-conformity score and prediction interval are derived using:

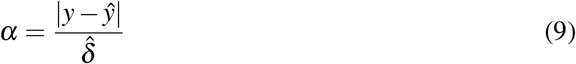

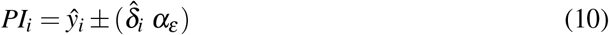

For a final independent and external evaluation of the OSSL models, we have tested several datasets depending on the spectral range of interest. In this complementary evaluation, partial least squares regression (PLSR) models were also fitted and optimized with 10-fold cross-validation (up to 30 components) for benchmarking performance, as this algorithm is a standard choice in the chemometrics field [55]. For the MIR range, the first test data set is composed of 70 samples scanned by several instruments (n = 20), which were obtained as part of the ring trial experiment developed by the SS4GG initiative [17]. This data set helps us to understand the robustness of OSSL models against small spectral variations resulting from dissimilarities across instruments and SOPs. Predictions models of `oc_usda.c729_w.pct`, `clay.tot_usda.a334_w.pct`, `ph.h2o_usda.a268_index`, and `k.ext_usda.a725_cmolc.kg` were investigated.

The second data set for MIR evaluation was compiled from long-term research (LTR) trial sites found throughout the US, spanning different locations and treatments such as cover crops, straw removal, fertilization, and irrigation [56]. This data set is especially attractive because it emulates a scenario of OSSL application and its effects for monitoring organic carbon changes across agricultural soils. This data set is publicly available with two spectral versions (KSSL and Woodwell, n = 162), where only `oc_usda.c729_w.pct` has reference analytical data.

For the VisNIR range, we also used the ring trial samples, although with a smaller sample size (n = 60) because only this amount was prepared as fine earth (soil particle size *<*2 mm) and shipped around the world. The same soil properties were evaluated using VisNIR, i.e., `oc_usda.c729_w.pct, clay.tot_usda.a334_w.pct`, `ph.h2o_usda.a268_index`, and `k.ext_usda.a725_cmolc.kg`. Lastly, the NIR Neospectra prediction models were tested with 90 soil samples from Ghana, Kenya, and Nigeria available through the public Neospectra database [32]. These samples were not used during model calibration but rather kept separated for external validation. Prediction models of `oc_usda.c729_w.pct`, `clay.tot_usda.a334_w.pct, ph.h2o_usda.a268_index`, and `k.ext_usda.a725_cmolc.kg` were tested.

## Results

### OSSL resources

The OSSL has compiled and harmonized the spectra from eleven different SSLs (Table 1). This was an enormous effort to make soil spectroscopy data more accessible by sharing a single endpoint, where everyone can access and fetch global data specific to their needs. The OSSL Manual describes different methods for getting data (https://soilspectroscopy.github.io/ossl-manual/database-access.html), which includes compressed .CSV and serialized files (.qs format) hosted on Google Cloud Storage, data collections imported to a NoSQL MongoDB, and several application programming interfaces (API) endpoints for filtering and selecting data from the database. Similarly, a graphical user interface (GUI) denominated OSSL Explorer (https://explorer.soilspectroscopy.org/) was also designed for exploring the database; however, it contains only the data with available spatial coordinates, not the full database. In fact, from more than 135,000 entries in the OSSL, 87,707 samples have spatial coordinates linked to at least one spectral region and one soil property of interest, as described in Table 2. This explains why some geographical regions are more represented than others in specific spectral regions (Fig. 3), e.g., the VisNIR range is more represented and concentrated over the European countries. Nonetheless, for those samples missing precise coordinates in the OSSL, county centroid coordinates were provided especially for the VisNIR and MIR spectra from the KSSL. In any case, the references of the original data producers are provided in the OSSL Manual if a user is interested in getting data with differing criteria. It is also important to mention that another available resource is the OSSL Engine (https://engine.soilspectroscopy.org/), an estimation service where users can upload spectra collected on their instruments and get back predictions with the response, uncertainty, and representation flag.

**Fig 3.**
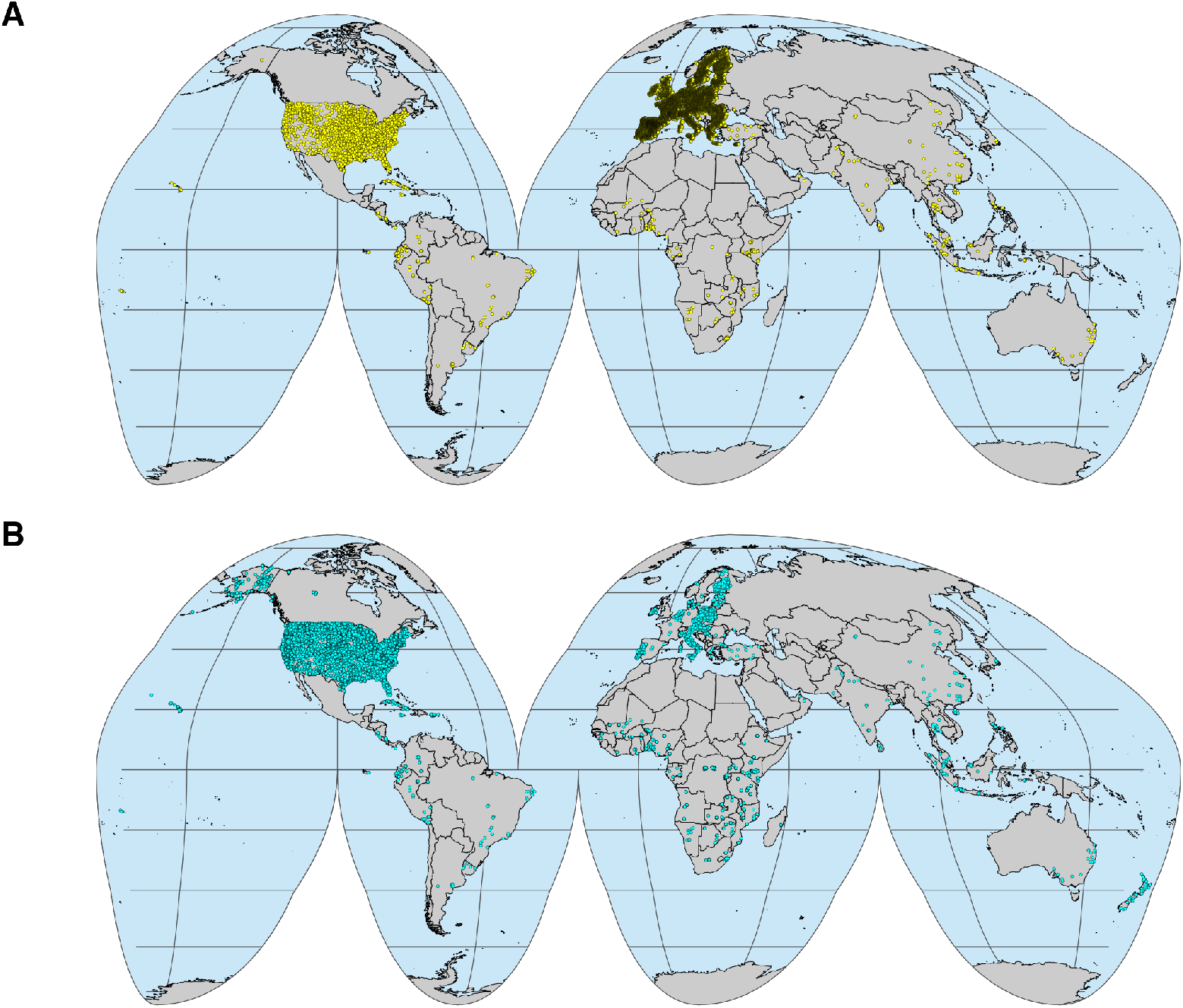
Geographical locations of the Open Soil Spectral Library (OSSL) samples. A: Locations with visible–near-infrared (VisNIR) spectra (350–2500 nm). B: Locations with mid-infrared (MIR) spectra (600–4000 cm^*−*1^). Note: not all OSSL samples have precise location data. From a total of 135,651 entries in the database, only 87,707 entries with varying complete information have precise location data. VisNIR samples in the US are plotted using county centroid coordinates.

### 10-fold cross-validation model performance

When OSSL was used to calibrate prediction models, the results for internal performance evaluation using 10-fold cross-validation (10CV) with refitting were highly variable depending on the soil property, the spectral region, and the subset of calibration (S1 Table). It is important to mention that we fitted separate models for the KSSL library to ensure that no systematic bias would be propagated to predictions due to the different analytical procedures used for defining the reference soil property values. This version of the model helped us to understand the effects of contrasting analytical methods on predictions and uncertainty, even after applying harmonization rules, when evaluating the larger models calibrated with the whole OSSL database.

As more than 140 models were fitted with the full OSSL database, we summarized internal evaluation performance by plotting Lin’s CCC against RPIQ values (Fig. 4). This plot shows the overall capacity of the models when linking the spectral variations to the original range of soil properties. Higher RPIQ values can be achieved either by having more broadly-distributed reference values (i.e., higher IQR) or by having more precise models, when the original calibration set is limited in representation (i.e., lower IQR). Both maximization, higher IQR or lower RMSE, results in higher RPIQ. Therefore, a cut-off of 2 means that the original variability measured by IQR is twice as large as the average model error (precision). It does not account for high bias (model inaccuracy), i.e., a potential mismatch caused by a shift of the predicted values even when getting precise models; thus Lin’s CCC enables us to account for this issue because it encompasses both the precision and accuracy of a model in its calculation.

**Fig 4.**
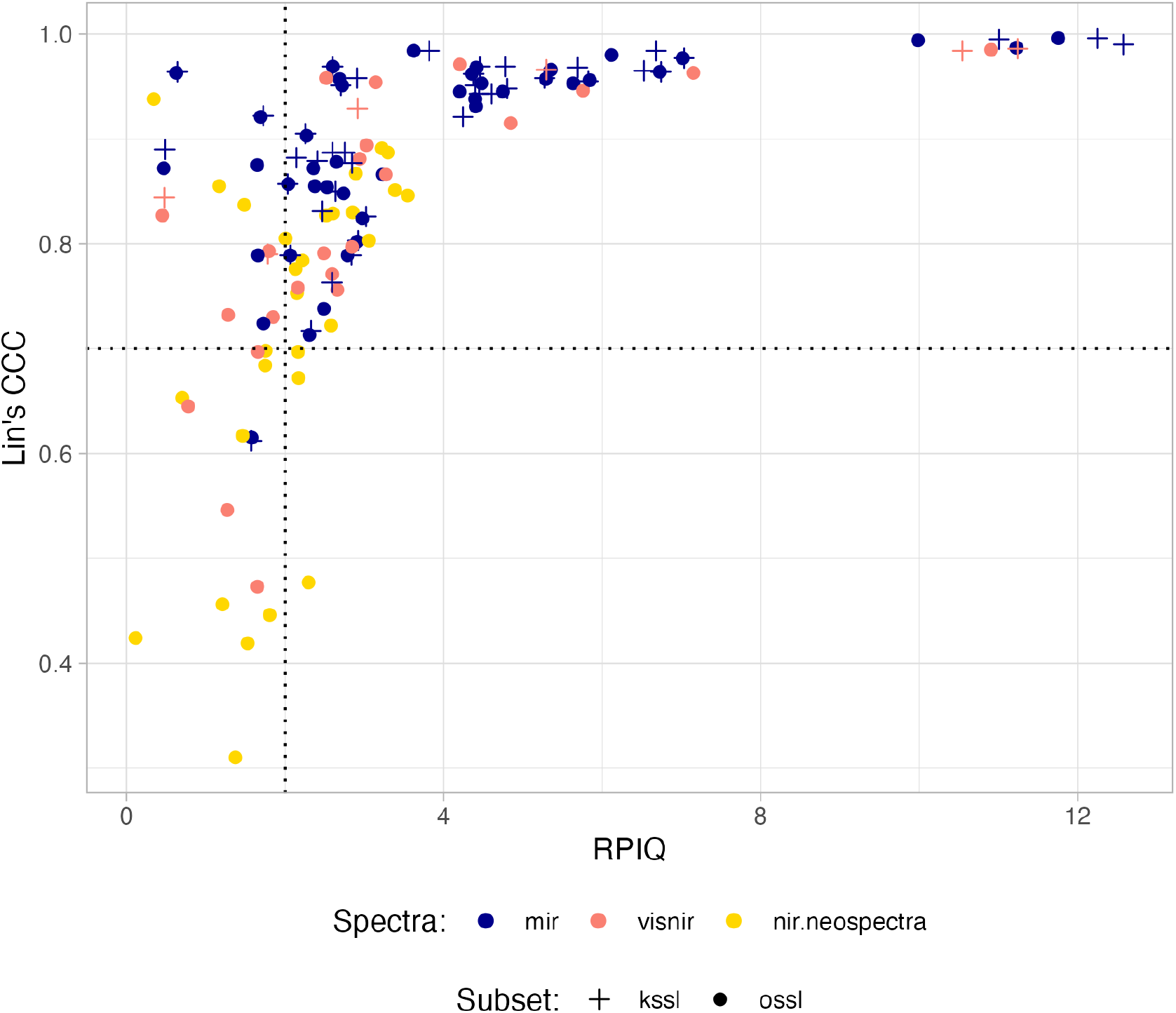
Internal performance (10-fold cross-validation) of prediction models fitted with the Open Soil Spectral Library (OSSL). Model performance among spectral ranges: Visible–near-infrared (350–2500 nm, visnir), mid-infrared (600–4000 cm^*−*1^, mir), and near-infrared (1350–2550 nm, nir.neospectra) specific to the Neospectra scanner (Si-Ware). Symbols reflect the models calibrated using the full OSSL database (ossl) or the Kellogg Soil Survey Laboratory alone (kssl). Note: Dashed lines are centered at Lin’s CCC of 0.7 and RPIQ of 2.

Setting thresholds for Lin’s CCC of 0.7 and RPIQ of 2, which convey both performance and extrapolation capacity, we observe that most of the models calibrated with the KSSL dataset were placed in the best performance quadrant (top right) (Fig. 4A). Similarly, the MIR range appears to be the spectral region with the best association with the variation in the original soil properties, followed by VisNIR and NIR (Neospectra), with NIR (Neospectra) having more limited models (bottom left quadrant, Fig. 4B). This classification indicates that it is important to use multiple goodness-of-fit metrics to account for the distinct types of errors (bias and variance) associated with the training capability (quantity and quality of samples) when assessing model performance.

Nonetheless, one can have an overview of the OSSL model’s calibration capacity ranked by soil property (Fig. 5). In this visualization, all the combinations of spectral regions for the whole OSSL are ranked in decreasing order by Lin’s CCC and striped when RPIQ ^*≤*^ 2. Overall, MIR had most of the soil properties’ models with Lin’s CCC *>* 0.7 and RPIQ *>* 2 when compared to VisNIR and Neospectra NIR. MIR models that did not pass this threshold classification were total sulfur (`s.tot_usda.a624_w.pct`), electrical conductivity (`ec_usda.a364_ds.m`), boron (Mehlich3 method, `b.ext_mel3_mg.kg`), sodium (both NH4OAc method `na.ext_usda.a726_cmolc.kg` and Mehlich3 method `na.ext_usda.a1068_mg.kg`), zinc (Mehlich3 method, `zn.ext_usda.a1073_mg.kg`), and available water content (`awc.33.1500kPa_usda.c80_w.frac`). For the VisNIR region, inconsistent calibration was found for bulk density (`bd_usda.a4_g.cm3`), coarse fragments (`cf_usda.c236_w.pct`), electrical conductivity (`ec_usda.a364_ds.m`), sodium (NH4OAc method `na.ext_usda.a726_cmolc.kg`), phosphorus (Olsen method, `p.ext_usda.a274_mg.kg`), total sulfur (`s.tot_usda.a624_w.pct`), water retention at 1500 kPa (`wr.1500kPa_usda.a417_w.pct`), and water retention at 33 kPa (`wr.33kPa_usda.a415_w.pct`). Lastly, for the Neospectra NIR models, calibration challenges were found for 12 models: Total aluminum (Crystalline form, `al.dith_usda.a65_w.pct`), total aluminum (Amorphous form, `al.ox_usda.a59_w.pct`), available water content (`awc.33.1500kPa_usda.c80_w.frac`), bulk density (`bd_usda.a4_g.cm3`), electrical conductivity (`ec_usda.a364_ds.m`), total iron (Amorphous form, `fe.ox_usda.a60_w.pct`), potassium (NH4OAc method, `k.ext_usda.a725_cmolc.kg`), manganese (KCl method, `mn.ext_usda.a70_mg.kg`), sodium (NH4OAc method, `na.ext_usda.a726_cmolc.kg`), phosphorus (Mehlich3 method, `p.ext_usda.a1070_mg.kg`), total sulfur (`s.tot_usda.a624_w.pct`), and water retention at 33 kPa (`wr.33kPa_usda.a415_w.pct`). This indicates that some regions have more limitations than others when only the spectral variations are used as predictors in the models. It is also clear that for many chemical soil properties, the calibration accuracy drops significantly from MIR to VisNIR and NIR instruments.

**Fig 5.**
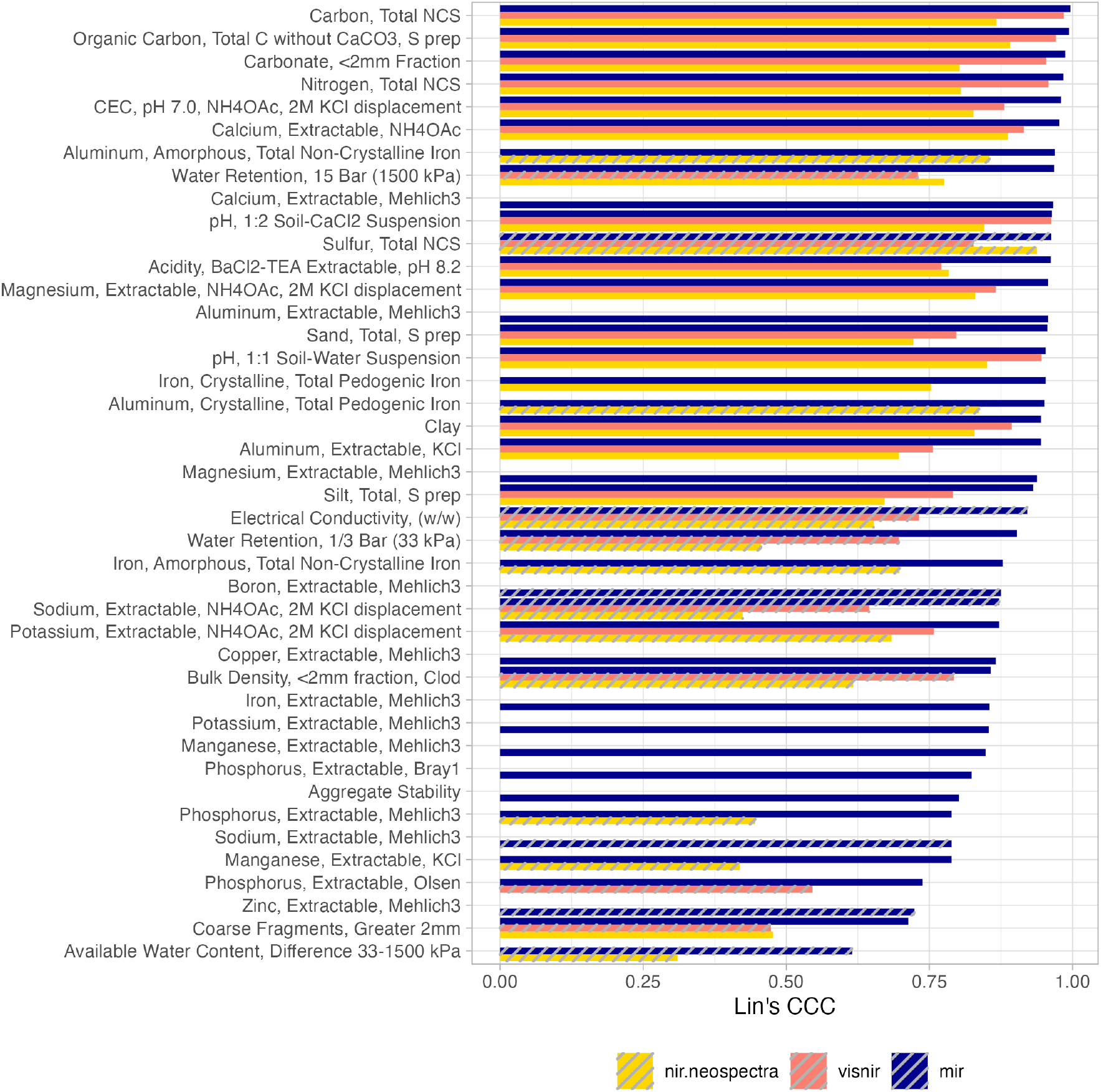
Rank of prediction models fitted with the Open Soil Spectral Library (OSSL). Lin’s concordance correlation coefficient (CCC) from 10-fold cross-validation with refitting. Striped bars represent models with a ratio of performance to the interquartile range (RPIQ) *<*2.

Users interested in the graphical analysis of a specific combination of soil property, spectral region of interest, and calibration library (either KSSL or the full OSSL), can have access to hex-binned scatter plots produced during model evaluation (Fig. 6). This type of visualization is included in the `ossl-models` GitHub repository and readily available via the OSSL Engine when selecting a model of interest.

**Fig 6.**
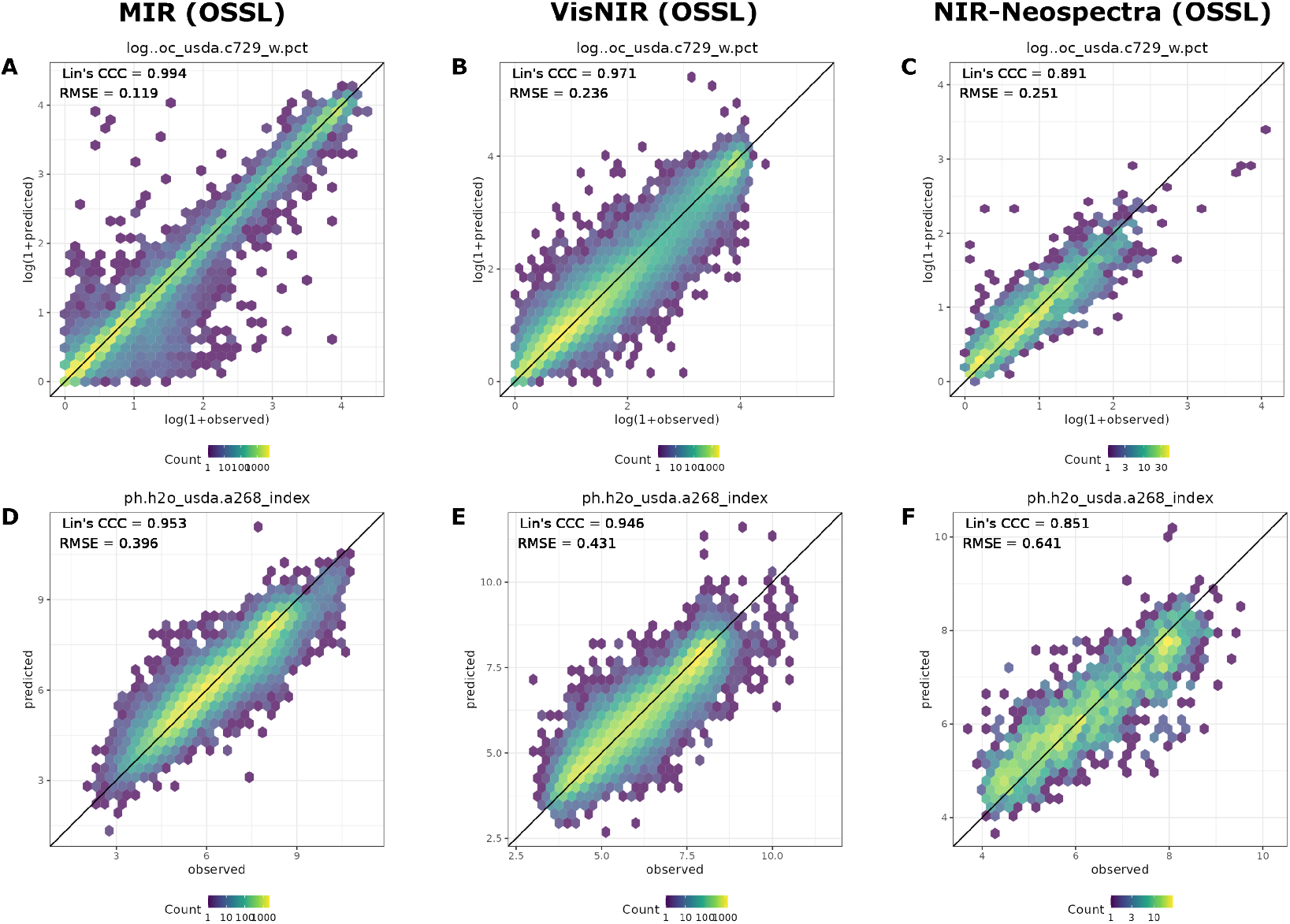
Example of accuracy plots for soil organic carbon (oc usda.c729 w.pct) content and soil pH in water (ph.h2o usda.a268 index) with Lin’s CCC gradually decreasing from MIR to NIR Neospectra models. A, B, and C are the accuracy plots of oc_usda.c729 w.pct from MIR, VisNIR, and Neospectra NIR models, respectively; D, E, and F are the accuracy plots for ph.h2o_usda.a268_index, respectively. All models have been fitted using Cubist. Accuracy results are based on 10–fold cross-validation with refitting.

### Independent evaluation

Using the VisNIR ring trial test set for the evaluation of OSSL models revealed a big variation caused by the origin of the spectra (Table 4). When we calculate the average (mean) for each model type, PLSR achieves a higher Lin’s CCC (0.67) and RPIQ (1.15) than Cubist (0.60 and 1.05, respectively). The performance edge from PLSR seems to be consistent across soil properties and the ossl subset, especially for `ph.h2o_usda.a268_index` (PLSR average Lin’s CCC of 0.74 *>* Cubist average Lin’s CCC of 0.65) and `k.ext_usda.a725_cmolc.kg` (PLSR average Lin’s CCC of 0.47 *>* Cubist average Lin’s CCC of 0.28). For other soil properties, the results were less variable. In fact, considering that the VisNIR ring trial is composed of twelve different instruments that measured the same soil samples, most of the variation is due to the differences among them, laboratory conditions, and procedures. For example, `clay.tot_usda.a334_w.pct`, `k.ext_usda.a725_cmolc.kg`, and `ph.h2o_usda.a268_index` were the most sensitive soil properties to the test spectral composition (see the minimum statistics of Lin’s CCC and RPIQ), but PLSR found a way to deliver slightly better predictions for those contrasting setups. Although the overall results were not satisfactory in some cases, the independent evaluation showed that some instruments have a spectral composition similar enough to the OSSL library and can be used with the OSSL Cubist models (hosted in the OSSL Engine) with satisfactory performance for `oc_usda.c729_w.pct`, `ph.h2o_usda.a268_index`, and `clay.tot_usda.a334_w.pct`, with superior performance found when using the KSSL subset. Nonetheless, even with a good indication of performance on these test instruments, new samples to be predicted must be properly represented by the calibration feature space to avoid prediction bias.

**Table 4.** Goodness-of-fit metrics from an independent evaluation of the visible–near-infrared (VisNIR) models calibrated with the Open Soil Spectral Library (OSSL). Statistics are reported for a sample of 12 distinct VisNIR instruments.

| Model type | Soil property | Model subset | Statistics | RMSE | bias | R<sup>2</sup> | CCC | RPIQ |
| --- | --- | --- | --- | --- | --- | --- | --- | --- |
| cubist | clay.tot.usda.a334_w.pct | ossl | min | 6.41 | -21.36 | 0.00 | -0.10 | 0.56 |
| cubist | clay.tot.usda.a334_w.pct | ossl | mean | 13.35 | -2.40 | 0.25 | 0.38 | 1.35 |
| cubist | clay.tot.usda.a334_w.pct | ossl | max | 27.96 | 4.40 | 0.70 | 0.80 | 2.46 |
| cubist | k.ext.usda.a725_cmolc.kg | ossl | min | 0.31 | -2.06 | 0.01 | 0.01 | 0.07 |
| cubist | k.ext.usda.a725_cmolc.kg | ossl | mean | 1.09 | -0.49 | 0.23 | 0.28 | 0.51 |
| cubist | k.ext.usda.a725_cmolc.kg | ossl | max | 4.41 | 0.16 | 0.66 | 0.47 | 1.00 |
| cubist | oc.usda.c729_w.pct | kssl | min | 1.63 | -4.31 | 0.76 | 0.63 | 0.19 |
| cubist | oc.usda.c729_w.pct | kssl | mean | 3.56 | -0.55 | 0.88 | 0.88 | 0.70 |
| cubist | oc.usda.c729_w.pct | kssl | max | 9.98 | 1.09 | 0.94 | 0.97 | 1.16 |
| cubist | oc.usda.c729_w.pct | ossl | min | 1.75 | -3.89 | 0.22 | 0.23 | 0.08 |
| cubist | oc.usda.c729_w.pct | ossl | mean | 5.27 | -0.88 | 0.78 | 0.82 | 0.65 |
| cubist | oc.usda.c729_w.pct | ossl | max | 22.78 | 0.36 | 0.94 | 0.97 | 1.07 |
| cubist | ph.h2o.usda.a268_index | ossl | min | 0.73 | -1.64 | 0.20 | 0.36 | 1.04 |
| cubist | ph.h2o.usda.a268_index | ossl | mean | 1.22 | -0.21 | 0.50 | 0.65 | 2.18 |
| cubist | ph.h2o.usda.a268_index | ossl | max | 2.25 | 0.53 | 0.85 | 0.84 | 3.20 |
| plsr | clay.tot.usda.a334_w.pct | ossl | min | 9.80 | -6.57 | 0.08 | 0.19 | 0.57 |
| plsr | clay.tot.usda.a334_w.pct | ossl | mean | 14.92 | 4.00 | 0.22 | 0.38 | 1.13 |
| plsr | clay.tot.usda.a334_w.pct | ossl | max | 27.77 | 14.53 | 0.36 | 0.59 | 1.61 |
| plsr | k.ext.usda.a725_cmolc.kg | ossl | min | 0.28 | -0.57 | 0.13 | 0.24 | 0.42 |
| plsr | k.ext.usda.a725_cmolc.kg | ossl | mean | 0.47 | -0.08 | 0.35 | 0.47 | 0.72 |
| plsr | k.ext.usda.a725_cmolc.kg | ossl | max | 0.75 | 0.37 | 0.48 | 0.68 | 1.12 |
| plsr | oc.usda.c729_w.pct | kssl | min | 1.72 | -5.18 | 0.68 | 0.61 | 0.20 |
| plsr | oc.usda.c729_w.pct | kssl | mean | 3.16 | -0.42 | 0.89 | 0.88 | 0.71 |
| plsr | oc.usda.c729_w.pct | kssl | max | 9.23 | 1.24 | 0.93 | 0.96 | 1.10 |
| plsr | oc.usda.c729_w.pct | ossl | min | 2.17 | -0.86 | 0.75 | 0.78 | 0.54 |
| plsr | oc.usda.c729_w.pct | ossl | mean | 2.65 | 0.57 | 0.92 | 0.89 | 0.73 |
| plsr | oc.usda.c729_w.pct | ossl | max | 3.52 | 1.47 | 0.96 | 0.93 | 0.87 |
| plsr | ph.h2o.usda.a268_index | ossl | min | 0.77 | -0.58 | 0.39 | 0.56 | 1.37 |
| plsr | ph.h2o.usda.a268_index | ossl | mean | 1.00 | -0.18 | 0.62 | 0.74 | 2.45 |
| plsr | ph.h2o.usda.a268_index | ossl | max | 1.71 | 0.25 | 0.79 | 0.83 | 3.06 |

RMSE: Root mean square error; bias: Mean error; R<sup>2</sup>: Coefficient of determination; CCC: Lin's concordance correlation coefficient; RPIQ: Ratio of performance to the interquartile range; plsr: Partial least square regression. Note: Metrics provided in original units after back-transformation.

In turn, the MIR ring trial test set employed for the evaluation of OSSL models revealed differing results that favor the use of Cubist models (Table 5). The average (mean) performance of the Cubist models was higher (Lin’s CCC of 0.79 and RPIQ of 2.19) than the PSLR models (Lin’s CCC of 0.76 and RPIQ of 1.87). Similarly, we found that models calibrated with the whole OSSL are, on average, slightly superior (Lin’s CCC of 0.78 and RPIQ of 2.11) than those fitted with the KSSL alone (Lin’s CCC of 0.77 and RPIQ of 1.96). Considering these favorable findings towards the use of Cubist in combination with the whole OSSL MIR spectra, we found that `c_usda.c729_w.pct` (Lin’s CCC and RPIQ of 0.95 and 2.96, respectively), `clay.tot_usda.a334_w.pct` (0.74 and 1.81), and `ph.h2o_usda.a268_index` (0.0.84 and 2.92) achieved satisfactory overall performance regardless of the instrument type. For `k.ext_usda.a725_cmolc.kg`, the performance was more limited (Lin’s CCC and RPIQ of 0.91 and 1.48, respectively). In any case, some instruments are still subject to unsatisfactory results, as they seem to have unusual spectral variations that cause poor performance.

**Table 5.** Goodness-of-fit metrics from an independent evaluation of the mid-infrared (MIR) models calibrated with the Open Soil Spectral Library (OSSL). Statistics are reported for a sample of 20 distinct MIR instruments. Metrics provided in original units after back-transformation.

| Model type | Soil property | Model subset | Statistics | RMSE | bias | R<sup>2</sup> | CCC | RPIQ |
| --- | --- | --- | --- | --- | --- | --- | --- | --- |
| cubist | clay.tot_usda.a334_w.pct | kssl | min | 4.22 | -37.80 | 0.19 | 0.09 | 0.36 |
| cubist | clay.tot_usda.a334_w.pct | kssl | mean | 8.76 | -1.60 | 0.70 | 0.79 | 2.14 |
| cubist | clay.tot_usda.a334_w.pct | kssl | max | 40.51 | 4.46 | 0.88 | 0.94 | 3.41 |
| cubist | clay.tot_usda.a334_w.pct | ossl | min | 5.62 | -8.42 | 0.01 | 0.06 | 0.78 |
| cubist | clay.tot_usda.a334_w.pct | ossl | mean | 8.82 | -1.84 | 0.62 | 0.74 | 1.81 |
| cubist | clay.tot_usda.a334_w.pct | ossl | max | 18.41 | 2.40 | 0.81 | 0.90 | 2.56 |
| cubist | k.ext_usda.a725_cmolc.kg | kssl | min | 0.24 | -3.12 | 0.00 | -0.00 | 0.12 |
| cubist | k.ext_usda.a725_cmolc.kg | kssl | mean | 0.70 | -0.23 | 0.45 | 0.60 | 1.26 |
| cubist | k.ext_usda.a725_cmolc.kg | kssl | max | 4.45 | 0.11 | 0.74 | 0.83 | 2.11 |
| cubist | k.ext_usda.a725_cmolc.kg | ossl | min | 0.22 | -1.17 | 0.04 | -0.13 | 0.30 |
| cubist | k.ext_usda.a725_cmolc.kg | ossl | mean | 0.44 | -0.07 | 0.49 | 0.61 | 1.48 |
| cubist | k.ext_usda.a725_cmolc.kg | ossl | max | 1.71 | 0.19 | 0.76 | 0.86 | 2.32 |
| cubist | oc_usda.c729_w.pct | kssl | min | 0.37 | -1.56 | 0.46 | 0.66 | 0.33 |
| cubist | oc_usda.c729_w.pct | kssl | mean | 1.38 | -0.26 | 0.94 | 0.96 | 2.10 |
| cubist | oc_usda.c729_w.pct | kssl | max | 5.79 | 0.47 | 1.00 | 1.00 | 5.11 |
| cubist | oc_usda.c729_w.pct | ossl | min | 0.36 | -1.53 | 0.62 | 0.32 | 0.33 |
| cubist | oc_usda.c729_w.pct | ossl | mean | 1.16 | -0.01 | 0.96 | 0.95 | 2.96 |
| cubist | oc_usda.c729_w.pct | ossl | max | 5.67 | 1.48 | 1.00 | 1.00 | 5.27 |
| cubist | ph.h2o_usda.a268_index | kssl | min | 0.41 | -0.09 | 0.13 | 0.13 | 0.93 |
| cubist | ph.h2o_usda.a268_index | kssl | mean | 0.81 | 0.37 | 0.76 | 0.81 | 2.84 |
| cubist | ph.h2o_usda.a268_index | kssl | max | 2.07 | 1.70 | 0.90 | 0.95 | 4.72 |
| cubist | ph.h2o_usda.a268_index | ossl | min | 0.43 | -0.88 | 0.39 | 0.49 | 1.41 |
| cubist | ph.h2o_usda.a268_index | ossl | mean | 0.72 | 0.11 | 0.75 | 0.84 | 2.92 |
| cubist | ph.h2o_usda.a268_index | ossl | max | 1.37 | 0.70 | 0.88 | 0.94 | 4.43 |
| plsr | clay.tot_usda.a334_w.pct | kssl | min | 5.86 | -12.51 | 0.04 | 0.15 | 0.65 |
| plsr | clay.tot_usda.a334_w.pct | kssl | mean | 9.94 | -1.63 | 0.57 | 0.71 | 1.64 |
| plsr | clay.tot_usda.a334_w.pct | kssl | max | 22.05 | 3.28 | 0.81 | 0.89 | 2.46 |
| plsr | clay.tot_usda.a334_w.pct | ossl | min | 5.29 | -12.58 | 0.07 | 0.22 | 0.70 |
| plsr | clay.tot_usda.a334_w.pct | ossl | mean | 9.11 | -2.13 | 0.65 | 0.74 | 1.82 |
| plsr | clay.tot_usda.a334_w.pct | ossl | max | 20.71 | 9.31 | 0.84 | 0.91 | 2.73 |
| plsr | k.ext_usda.a725_cmolc.kg | kssl | min | 0.30 | -0.21 | 0.00 | 0.00 | 0.35 |
| plsr | k.ext_usda.a725_cmolc.kg | kssl | mean | 0.44 | 0.14 | 0.43 | 0.58 | 1.35 |
| plsr | k.ext_usda.a725_cmolc.kg | kssl | max | 1.46 | 1.39 | 0.56 | 0.71 | 1.72 |
| plsr | k.ext_usda.a725_cmolc.kg | ossl | min | 0.29 | -0.39 | 0.01 | 0.01 | 0.44 |
| plsr | k.ext_usda.a725_cmolc.kg | ossl | mean | 0.41 | 0.04 | 0.45 | 0.60 | 1.42 |
| plsr | k.ext_usda.a725_cmolc.kg | ossl | max | 1.16 | 1.05 | 0.58 | 0.72 | 1.74 |
| plsr | oc_usda.c729_w.pct | kssl | min | 0.51 | -1.80 | 0.88 | 0.17 | 0.26 |
| plsr | oc_usda.c729_w.pct | kssl | mean | 1.60 | 0.13 | 0.98 | 0.93 | 2.02 |
| plsr | oc_usda.c729_w.pct | kssl | max | 7.14 | 2.69 | 0.99 | 1.00 | 3.70 |
| plsr | oc_usda.c729_w.pct | ossl | min | 0.63 | -1.64 | 0.86 | 0.24 | 0.29 |
| plsr | oc_usda.c729_w.pct | ossl | mean | 1.59 | 0.14 | 0.98 | 0.93 | 1.85 |
| plsr | oc_usda.c729_w.pct | ossl | max | 6.49 | 2.24 | 0.99 | 0.99 | 3.00 |
| plsr | ph.h2o_usda.a268_index | kssl | min | 0.60 | -0.45 | 0.25 | 0.24 | 0.59 |
| plsr | ph.h2o_usda.a268_index | kssl | mean | 1.00 | 0.26 | 0.75 | 0.79 | 2.28 |
| plsr | ph.h2o_usda.a268_index | kssl | max | 3.24 | 2.22 | 0.83 | 0.89 | 3.21 |
| plsr | ph.h2o_usda.a268_index | ossl | min | 0.56 | -0.35 | 0.47 | 0.56 | 1.22 |
| plsr | ph.h2o_usda.a268_index | ossl | mean | 0.81 | 0.12 | 0.78 | 0.83 | 2.59 |
| plsr | ph.h2o_usda.a268_index | ossl | max | 1.58 | 1.20 | 0.84 | 0.90 | 3.44 |

The previous findings were also confirmed by another evaluation (Fig. 7). The combination of the Cubist +?OSSL subset resulted in a lower error (RMSE) for the estimation of organic carbon for agricultural experimental sites, regardless of the origin of the spectra (KSSL or Woodwell spectra). The KSSL subset was only comparable when the test spectra were collected using the same instrument. In addition, the PLSR RMSE was always higher than the Cubist for all cases. Additional metrics for MIR data from LTR sites are provided in S2 Table.

**Fig 7.**
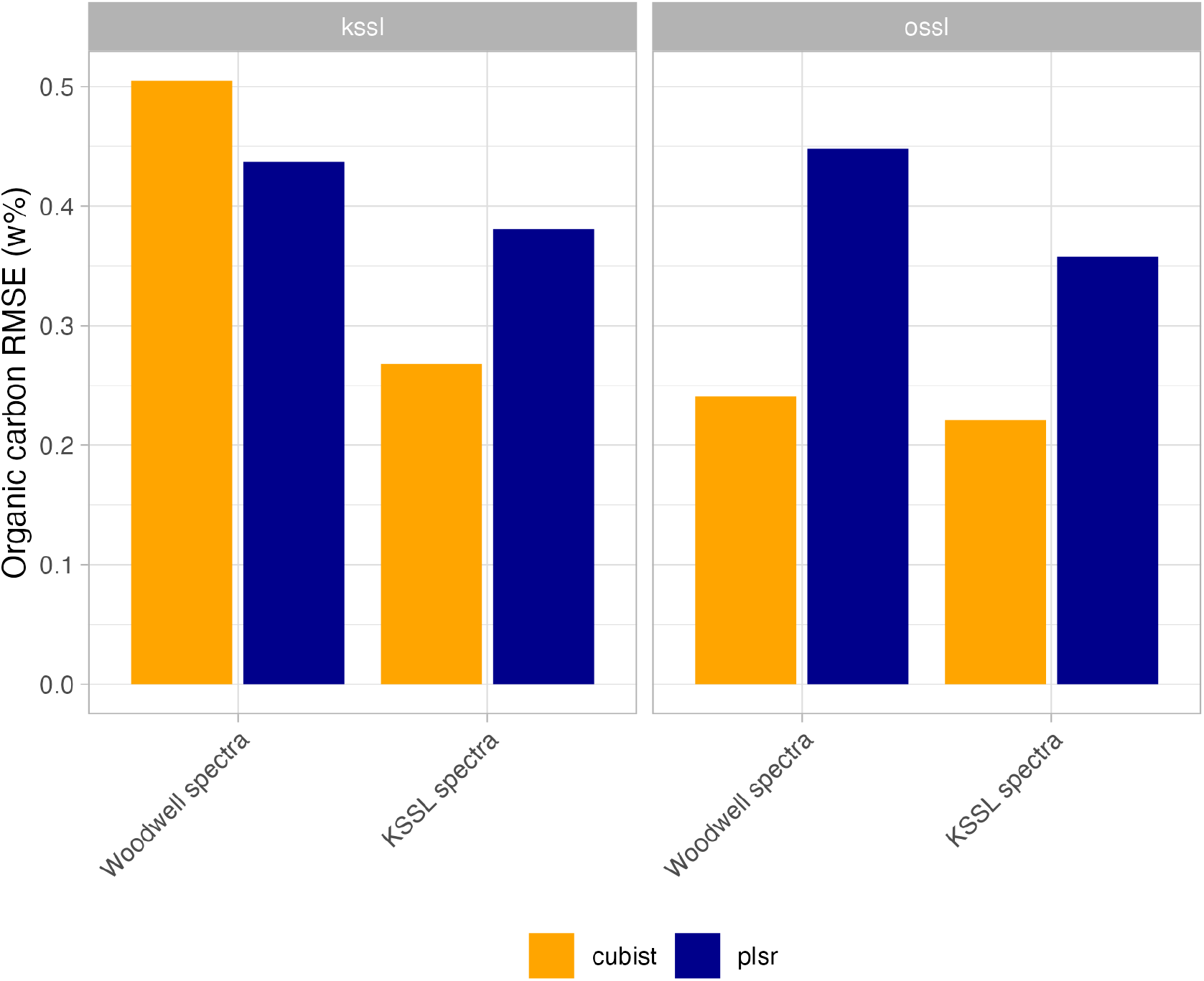
Root Mean Square Error (RMSE) of organic carbon from long-term research (LTR) sites predicted with the Open Soil Spectral Library (OSSL) models. ossl: models calibrated with the full OSSL. kssl: models calibrated with the Kellog Soil Survey Laboratory data alone. Note: Two different spectra versions are tested (n=162), i.e. scanned both at Woodwell Climate and KSSL.

Lastly, the results of the independent evaluation of the OSSL models for the NIR (Neospectra) range using the African soil samples (Table 6) revealed that Cubist models are again generally better (mean Lin’s CCC and RPIQ of 0.55 and 1.40, respectively) than the baseline models fitted with PLSR (mean Lin’s CCC and RPIQ of 0.49 and 1.22, respectively). For the Cubist models, the best performance was found for `oc_usda.c729_w.pct` (Lin’s CCC and RPIQ of 0.85 and 1.53, respectively), while `k.ext_usda.a725_cmolc.kg` only achieved Lin’s CCC and RPIQ of 0.43 and 1.01, respectively. In fact, PLSR was only superior for this soil property. The results of the Neospectra independent validation show that, for this limited spectral range, the performance is inferior to other spectral ranges (VisNIR and MIR) and more generally useful only for a limited number of soil properties. In addition, the contrasting geographical origin between the training and test samples also may pose limitations on the true verification of the Neospectra models. When they were tested against the calibration feature space of the OSSL (US-only samples), all the African soil samples were flagged as represented by the spectral variation (based on the Q-statistics method).

**Table 6.** Goodness-of-fit metrics from an independent evaluation of near-infrared (Neospectra) models calibrated with the Open Soil Spectral Library (OSSL).

| Model type | Soil property | RMSE | bias | R<sup>2</sup> | CCC | RPIQ |
| --- | --- | --- | --- | --- | --- | --- |
| cubist | clay.tot_usda.a334_w.pct | 14.21 | 2.77 | 0.47 | 0.59 | 1.72 |
| cubist | k.ext_usda.a725_cmolc.kg | 0.30 | 0.02 | 0.20 | 0.43 | 1.01 |
| cubist | oc_usda.c729_w.pct | 0.46 | 0.09 | 0.84 | 0.85 | 1.53 |
| cubist | ph.h2o_usda.a268_index | 1.06 | 0.09 | 0.11 | 0.32 | 1.35 |
| plsr | clay.tot_usda.a334_w.pct | 16.84 | -0.53 | 0.25 | 0.46 | 1.45 |
| plsr | k.ext_usda.a725_cmolc.kg | 0.31 | -0.08 | 0.24 | 0.47 | 0.96 |
| plsr | oc_usda.c729_w.pct | 0.51 | -0.13 | 0.75 | 0.84 | 1.39 |
| plsr | ph.h2o_usda.a268_index | 1.30 | 0.45 | 0.04 | 0.19 | 1.10 |

RMSE: Root mean square error; bias: mean error; R<sup>2</sup>: Coefficient of determination; CCC: Lin's concordance correlation coefficient; RPIQ: Ratio of performance to the interquartile range; plsr: Partial least square regression. Note: Metrics provided in original units after back-transformation.

## Discussion

### Summary findings

The OSSL is formed by different data sets with variable numbers of soil properties and spectral ranges. This resulted in the calibration of 141 models with varying internal and external performance (Fig.4). However, it is essential to consider that for the first time, a global spectral library is freely available under an open data license that covers nearly all continents. This geographical representation is unprecedented since the first use of a spectrometer to quantify soil properties [57]. Each model was fitted on the basis of the specificity of the spectral range, spectrometer’s manufacturer (i.e., the NIR Neospectra models), two different calibration libraries (KSSL alone or full OSSL), wet chemistry methodology (some chemical soil properties have distinct methods available), and so on. Therefore, users can select the configuration that best fits their needs. While preconfigured models are provided, more advanced users can access the data and build any type of model that will suit their needs.

On average, the MIR range appears to be the best spectral region for developing spectral prediction models, followed by VisNIR and NIR (Neospectra), in which the latter can provide good external performance for `oc_usda.c729_w.pct` (i.e., soil organic carbon) and `clay.tot_usda.a334_w.pct` (i.e., clay content) given its limited spectral coverage and cost. This happens because the MIR contains several fundamental and resolved absorption features from mineral and organic functional groups that translate to better prediction capacity, despite challenges in the interpretation that stem from chemical heterogeneity [58]. VisNIR and NIR spectra, in turn, are made of overtones from the fundamental vibrations of the MIR range, hence, are less sensitive to soil constituents and may result in inferior performance. In addition, we found good performance for some soil properties that may not directly affect soil spectra but can be indirectly inferred and quantified (secondary properties), such as cation exchange capacity, pH, soil contaminants, etc. However, understanding both primary and secondary components in the soil helps to better understand the factors that contribute to the improvement of spectral predictive models within the complex context of soil systems and also to select the spectral range, where they are most pronounced. On average, the performance results of the OSSL models are consistent with findings documented in the literature [14, 16, 59, 60].

We also found that the external performance (independent evaluation) is highly dependent on the characteristics of the test spectra, like spectral composition influenced by the instrument and standard operating procedures (SOPs). Using the experimental data from a ring trial [17], we found that VisNIR was more sensitive to instrument dissimilarity, with PLSR being the best-performing prediction algorithm. For the MIR and NIR (Neospectra) ranges, the Cubist models showed superior performance and were favored by the integration of several original SSLs that composed the OSSL. Nevertheless, we cannot exclude the effect of the origin and quality of the original datasets that make up the VisNIR calibration set. This yields variable results and the use of good SOPs with quality assurance and quality control (QA/QC) is an essential part of soil spectroscopy, regardless of the spectral range measured [11, 17].

### Reliability of predictions

Building trust in predictive modeling requires the provision of complementary information, as the predictions might still be biased and/or with high variance even from satisfactory calibrated models. Going beyond the standard evaluation of models using goodness-of-fit metrics, the OSSL provides the precision of each prediction using the conformal prediction method (Fig. 8). Different model types (Cubist and PLSR) resulted in different prediction intervals for soil organic carbon (SOC, `oc_usda.c729_w.pct`) when testing the LTR soil samples. Cubist models had a median width interval of 0.6 and 0.88, depending on the origin of the spectra, while PLSR resulted in broader median widths of 1.54 and 1.83. The assessment of all combinations (model type and spectra source) indicated coverage of the true values around or higher than the confidence level (95%, as the error probability was set to 5%), i.e., no model failed to translate its accuracy and precision to the predicted intervals. However, Cubist was more consistent than PLSR when combined with conformal prediction, as it resulted in the best goodness-of-fit model metrics coupled with a more approximate coverage to the predefined error and narrower uncertainty intervals.

**Fig 8.**
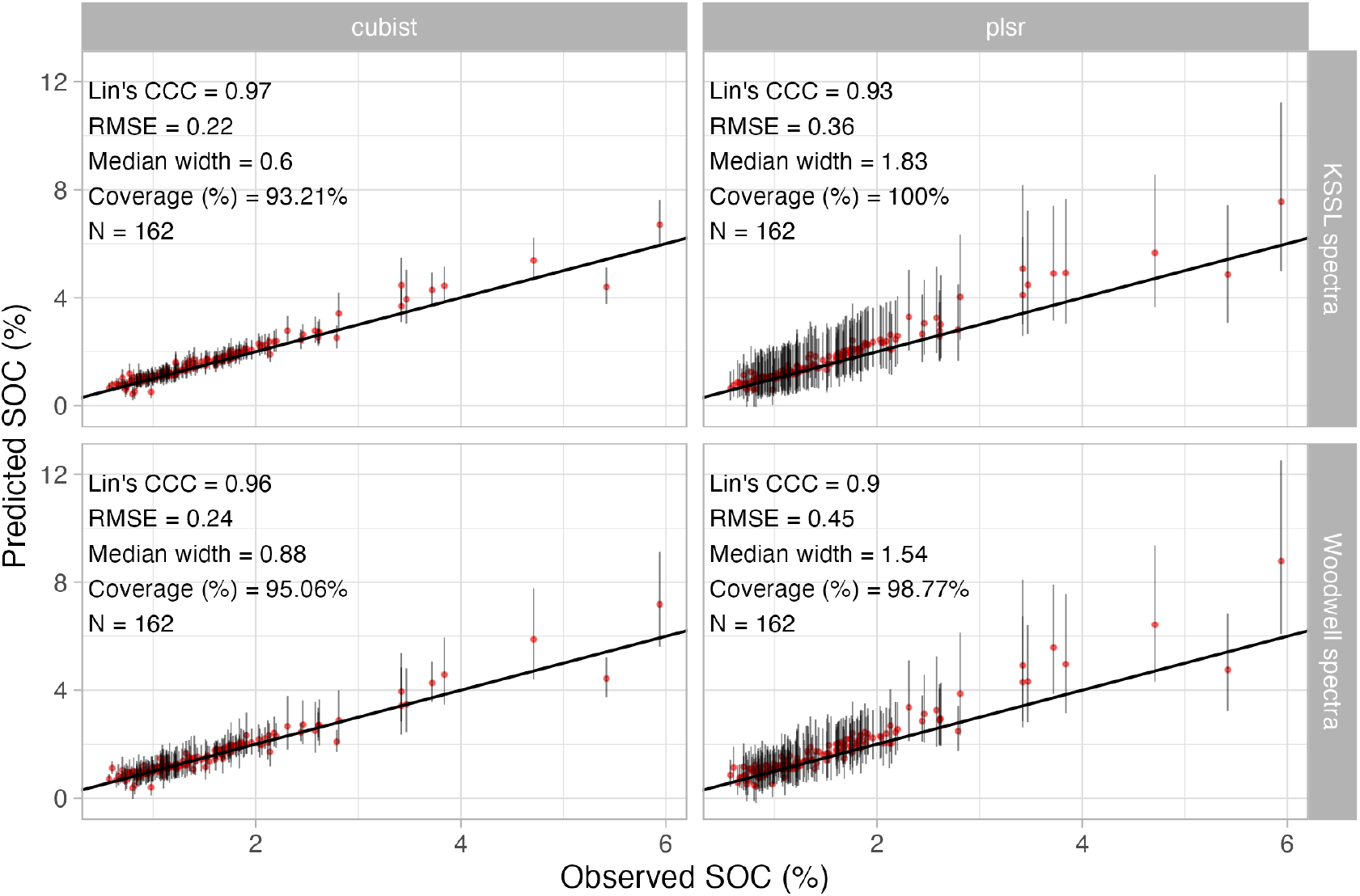
Uncertainty of predicted soil organic carbon (SOC, oc usda.c729 w.pct) values (precision at 5% error probability) from models fitted with the whole Open Soil Spectral Library database. The left and right panels depict the uncertainty of Cubist and Partial Least Square Regression (PLSR) models, respectively. The top and bottom panels depict the uncertainty from the long-term research (LTR) sites scanned both in the Kellog Soil Survey Laboratory and Woodwell Climate, respectively. Note: all metrics are provided in the original units after backtransformation. The median width and coverage of the prediction intervals (PI95%) are provided for the uncertainty assessment.

Another element of the reliability assessment is the screening of potential outliers or underrepresented samples regarding the calibration feature space. This mechanism may help to understand the capacity of a model to extrapolate by simultaneously avoiding high bias on new predictions. There are different ways for this purpose but the soil spectroscopy literature has been promoting the use of process control charts that rely upon one or multiple quantitative methods [48, 61]. Q-statistics is routinely employed in control charts and has the advantage of leveraging compression or orthogonalization algorithms (like PCA and PLS) used in predictive modeling. However, the list of methods for process control is long, including Hotelling *T* ^2^ test, F-ratio, SIMCA, etc [44, 61]. Further research still needs to be carried out especially because most of these methods rely on assumptions like the multivariate normal distribution and may be sensitive to heterogeneous soil spectral libraries, like the OSSL. In addition, the current orthogonalization implemented in the OSSL considers the full spectral space rather than the subset of every single combination of a soil property with the spectral region of interest, potentially yielding suboptimal results.

In the OSSL, Q-statistics was implemented by comparing the difference between the compressed back-transformed spectra and its original version with a critical value from the training set (Fig. 9). In an illustration example, cropland soils from Massachusetts, USA, were flagged as underrepresented likely because they have distinct spectral patterns than the Neospectra calibration dataset. Not all soil types and geographical regions are properly represented in the calibration samples as it is a smaller representation of the KSSL soil survey database, so the Q-statistics was able to flag them based solely on spectral representation. This indicates that new soil sampling on underrepresented places or the design of new projects to specific needs (e.g., organic carbon change monitoring) will still be crucial to leverage the full potential of spectral prediction models without introducing prediction bias [8].

**Fig 9.**
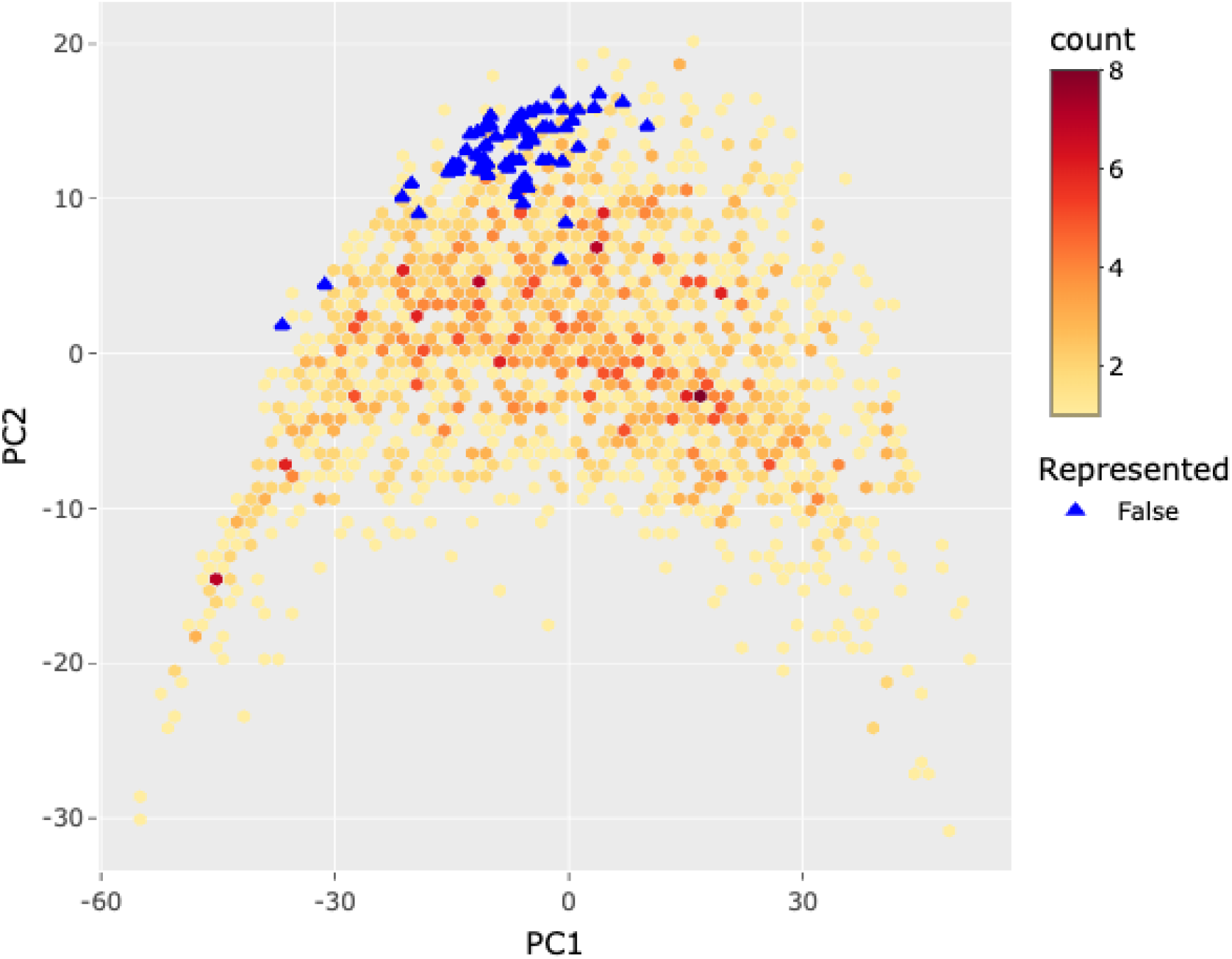
Example of the representation flag based on principal component analysis (PCA) and Q-statistics. In this example, samples from farmland soils (Massachusetts, USA) were evaluated against the NIR Neospectra OSSL model. All the test samples fell in the edge of the first two PCA space and were flagged as underrepresented regarding the full spectral feature space from the OSSL database. Note: The Q-statistics method uses all the retained principal components (in our study, 120 components) for flagging underrepresented spectra, not only the first two components depicted in the illustration. Figure sourced from the OSSL Engine (https://engine.soilspectroscopy.org/).

### An open and reproducible science initiative

The OSSL is a genuine open science project, meaning that it is a reproducible project aimed at the early sharing of soil data, research, and technological developments including open data, open standards, open source software, and open communication [62]. In fact, one of the main pillars of SS4GG is to support open innovation, so that all relevant knowledge actors are included and involved in the project by posting most of the input data sets, codes, and outputs on Github and similar repositories. Thus, all future exploitable activities of research outputs can be considered Findable, Accessible, Interoperable, and Reusable (FAIR). In this sense, OSSL can be considered as an open source driver for soil spectroscopy comparable to the widely used solutions for geospatial data, for example, the Geospatial Data Abstraction Library (GDAL) [63] and the Global Biodiversity Information Facility (GBIF) [64], to mention just a few projects that have especially inspired us.

There are limitations to fully open research especially regarding data privacy. One could argue what is more important for the global good: (A) the privacy and data ownership rights of farmers/land owners, or (B) the common interest of the public. Without proposing that either of the two is more important, we (the authors in this article) in fact support both paths, as the world’s soils are currently endangered [65, 66] and we need all possible efforts, open-source and commercial, to help conserve, regenerate, and improve soils in landscapes. In that sense, OSSL could potentially also host a number of proprietary datasets, provided that at least the derived models can be served as open data (Fig. 10). However, we hope that most of the legacy datasets will eventually be released under an open data license and be included in the OSSL, especially those that have already been funded by public money through taxpayers.

**Fig 10.**
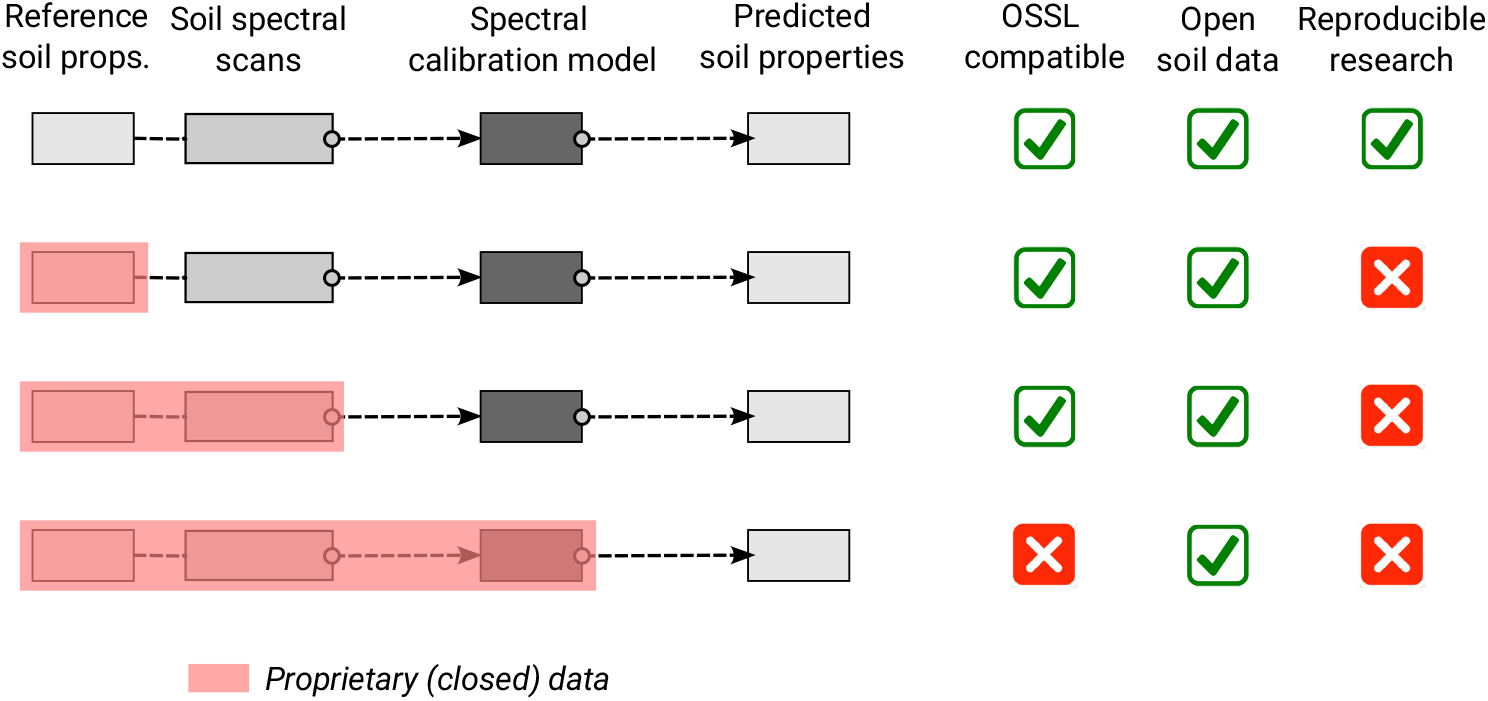
License matrix and possible combinations of open data and reproducibility. Many datasets can be included in OSSL; however, some minimum license compatibility is needed. OSSL accepts a diversity of soil spectral data sets as long as at least the produced models (model parameters) are shared under an open data license.

### Limitations and future development

We envision that soil spectroscopy in the near future will become more accessible, more affordable, and better integrated into the measurement, monitoring, reporting, and verification (MMRV) of soil status and ecosystem services. This is simply because soil monitoring is a growing market, and soil spectroscopy will inevitably play a role in that growth. The following three aspects of the development of soil spectroscopy seem to be especially promising: (1) miniaturization of field-scale spectrometers, (2) effective international integration for building global soil spectral libraries and standard operating procedures, and (3) continued research and development of learning algorithms for improving predictive capacity.

The miniaturization and cost reduction of handheld field spectrometers, especially in the NIR and MIR range, can revolutionize soil spectroscopy adoption and increase the number of applications multifold [67]. Several interesting handheld or portable spectral scanners are used today to generate thousands of soil scans, e.g. Neospectra™ [68, 69] and trinamiX™. Such light and portable NIR spectrometers could be used to take hundreds of georeferenced soil scans per day, improving the representation of variable soil landscapes. However, field soil spectroscopy has many challenges that are not common to laboratory soil spectroscopy, where conditions are more controlled. Soil moisture content, variable surface conditions, and environment conditions (for passive instruments) are just a few worth mentioning that have a large impact on the quality of the field spectra, and therefore, must be properly accounted for when integrating or working with dry-scan, laboratory-based soil spectral libraries, like the OSSL [70]. Nonetheless, with appropriate labeling and metadata compilation, the OSSL can accommodate different levels of spectral acquisition (e.g. laboratory and field) and enable adequate interoperability solutions.

In terms of international cooperation and integration, the IEEE AS P4005 working group has been working to define standards and protocols for laboratory, field, and remote sensing soil spectroscopy. FAO’s GLOSOLAN has also been defining the standard operating procedures (SOPs) for four categories of soil properties useful for international projects and extended soil health monitoring [71, 72], the same that were compiled in the OSSL depending on the availability of the original datasets. Likewise, GLOSOLAN has been leading the discussion and the definition of best practices for ensuring quality in the process of generating soil information via analytical wet chemistry. Not only the harmonization of analytical procedures is critical for soil data generation, but also the standardization across instruments and spectroscopy laboratories. A dedicated branch was created for dry chemistry (i.e. soil spectroscopy) in the GLOSOLAN network to promote and establish soil spectroscopy as a clean and sustainable solution for generating soil information across routine and research laboratories. In a parallel effort that was inspired by the previous, SS4GG has recently established a network with more than 20 participants across the world to assess dissimilarities across instruments and laboratories. The first assessment report has been published for the MIR range [17], but we hope that continued research and novel solutions keep coming to ensure better interoperability of soil spectroscopy laboratories and the OSSL.

For the third point on enhancing predictive performance, although the OSSL models showed promising results, several approaches can potentially provide more reliable results on any spectra uploaded to the OSSL. One improvement path would be incorporating ancillary information during model calibration upon the provision of coordinates, date, and depth for new samples. Many approaches can be employed to improve the prediction performance of spectral models as we have indicated that some spectral regions achieved limited prediction performance (Fig. 5). Multisensor and data fusion appear promising, as they support exploring complex interactions and variations of the landscape by the incorporation of spatial/temporal context in the models, minimizing in this case, spatial clustering or the overrepresentation of topsoil (Fig. 11). One limitation of such an approach, however, is that all the values of the covariates are needed for new samples, and hence this approach is heavily dependent on the availability of up-to-date data and user input of precise spatial and temporal sampling information. This, in fact, led to the simplification of the first OSSL models to rely just on spectral variations and specific ranges, as more sophisticated approaches would create barriers to end-user adoption.

**Fig 11.**
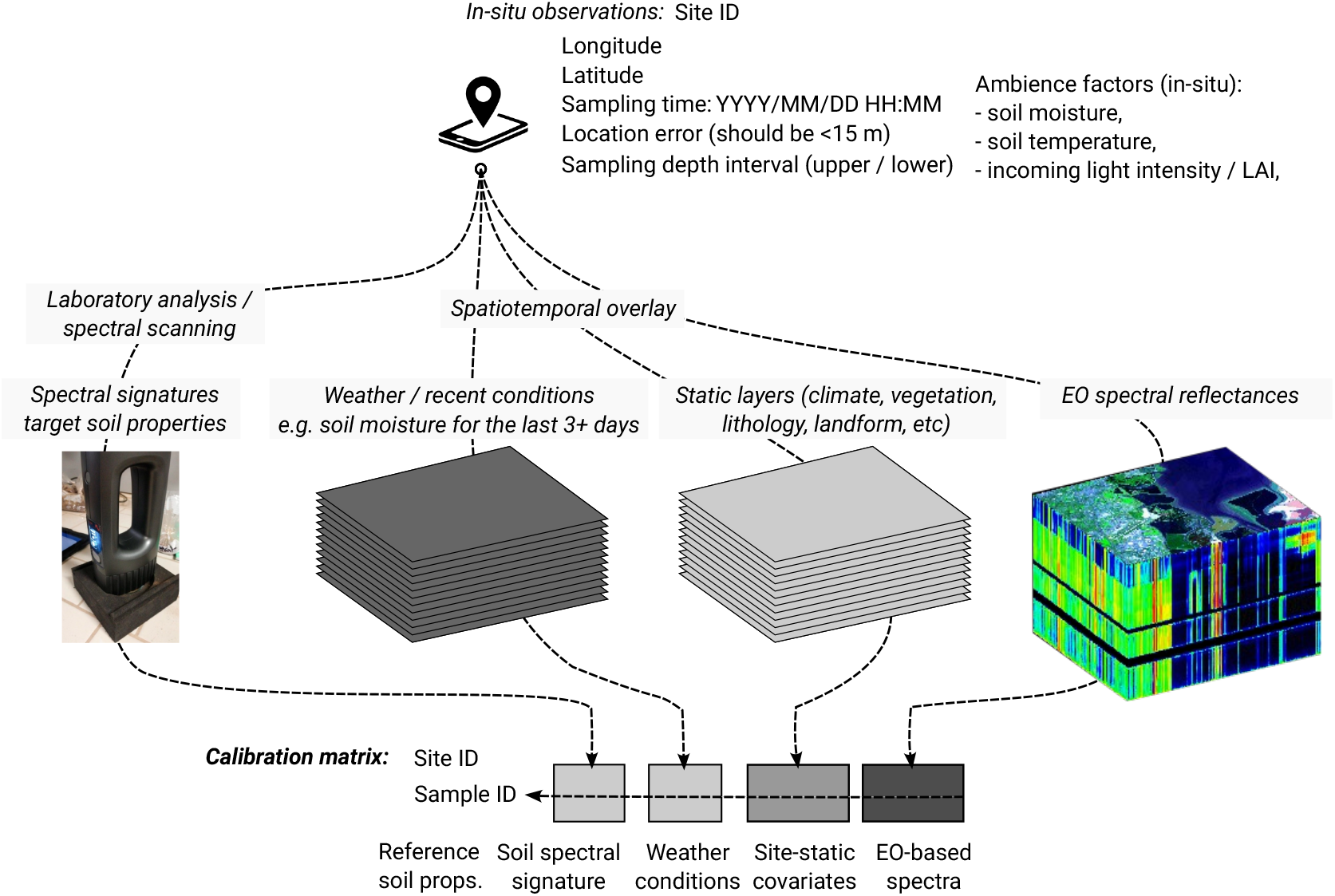
A data processing scheme for spatially explicit soil spectral calibration. Variables and data sources of interest for producing spatially explicit calibration models. This however requires that all soil samples are fully georeferenced and that covariate layers are available for new prediction locations (including forecasted predictions of weather conditions and similar). Portable NIR scanner in the picture: Neospectra Scanner covering the range from 1350 to 2500 nm.

Today, many detailed auxiliary spatial-temporal data are available, even with global coverage, that could be used to improve the calibration of *in situ* soil spectral libraries. For example, soil moisture often drastically changes spectral signatures that negatively impact the predictions, so incorporating a detailed estimate of daily soil moisture may help improve the accuracy of spectral models [73–75]. Spectral reflectance information from Earth Observation systems, e.g., VisNIR reflectances from Sentinel-2 or Landsat, could be explored with a special focus on bare/no-vegetation spectral reflectance of the land surface [76, 77]. Nonetheless, EO spectral data may have delays and yield additional costs, as it often takes a significant amount of time to process and prepare cloud-free, bare-earth, or similar high-quality products. Another critical factor about integrating EO reflectance products with field measurements is that, although there are several ground observation points with reference spectral data, the compatibility between the different platforms (field sensor, aerial, orbital) still seems to pose a big barrier and requires further research and development for proper integration.

Model localization is another approach that seems very promising in predictive soil spectroscopy but may pose some restrictions when building estimation services as they often require recursive processing and retraining of model parameters [78, 79]. In addition, local models can produce unreliable results if there is a low similarity between the local and reference library [80], indicating that compilation of large spectral libraries (like the OSSL) is critical for increasing the representation of variable soil types and conditions. Generally applicable models, like those employed in the OSSL, have less of this localization problem but might fail by delivering biased predictions on local samples. Cutting-edge machine learning methods, especially those under the umbrella of deep learning, could be further explored to get the best out of large datasets with multitask learning (prediction of multiple soil properties at the same time), transfer learning between ranges (VisNIR, NIR, and MIR) and domains (laboratory, field and remote), and also allowing the public storage and distribution of pre-trained large models. In fact, both deep learning and localization approaches have been recently demonstrated to work well when combined with the Neospectra NIR for predicting soil organic carbon (25% improvement over the current OSSL framework) [81]. This happened after the promotion of a community data science competition on Kaggle as part of the SS4GG event series, indicating an effective path for community-based model improvement and testing of new ideas with the OSSL.

### Please, contribute!

OSSL is a genuine open science project with (both) open-source and open-data licenses (https://opendefinition.org/licenses/). We aim to develop and improve calibration models through open science hackathons and workshops. All of our code can be found online under the MIT license; a versioned backup copy of the data is also available via Zenodo under the CC-BY license. Both licenses allow you to extend, build on, and even build commercial businesses on top of this data and code. The data we have created have already been used in some projects [82, 83]. Help us create better models for global good and especially to support nature-based solutions, regenerative agriculture, and environmental monitoring projects. If you are the owner of soil spectral libraries, consider donating your data to the project. Your data will be systematically imported into OSSL and used to update global and local soil spectroscopy calibration models; your attribution and citation requests will be carefully followed.

SS4GG has been encouraging researchers and entities (public and private) to support the project by sharing any SSL. We have been promoting four modes of data sharing: (i) publishing the SSL with an open access license by releasing it under a Creative Commons license (CC-BY, CC-BY-SA) or the Open Data Commons Open Database License (ODbL), so we can directly import into the OSSL with proper attribution; (ii) donating a small portion (e.g. 5–10%) of the dataset under CC-BY, CC-BY-SA and/or ODbL, which can also be directly imported into the OSSL with proper attribution; (iii) allowing the SS4GG initiative to directly access the data so that we can perform data mining and then release only the models under an open data license; and/or (iv) using the OSSL database to produce new derivative products, then sharing them through our own infrastructures or contact us for providing hosting support. In addition, we can sign professional data-sharing agreements with data producers that specify in detail how the data will be used. Our primary interest is in enabling research, sharing, and use of models (calibration and prediction) in collaboration with groups across borders.

## Conclusion

We have described a comprehensive and curated global soil spectral library called *“Open Soil Spectral Library”* (OSSL) which is based on open data principles for building a collaborative and inclusive community of soil spectroscopy practitioners and stakeholders. The OSSL is composed of a suite of datasets, reproducible code, web services, and tutorials. It is one of the main outputs of the Soil Spectroscopy for Global Good (SS4GG) project which aims at bridging gaps between new technology and wider use for soil monitoring. The results of model cross-validation (10–fold with refitting) for MIR and Vis-NIR spectra show that, in general, MIR-based models are more accurate than VisNIR-based models, especially for determining chemical soil properties. From an independent test evaluation, the Cubist comes out as the best-performing algorithm for global calibration and generation of additional reliable outputs, i.e., prediction uncertainty and representation flag. Although many soil properties are well predicted, some properties such as total sulfur, extractable sodium, and electrical conductivity, performed poorly in all spectral regions, with some other extractable nutrients and physical soil properties (coarse fragments, soil water retention parameters, and bulk density) also performing inadequately in one or two spectral regions (VisNIR or Neospectra NIR). Hence, the use of predictive models based solely on spectral variations has limitations. With this genuinely open-source and open-access project, we hope that OSSL becomes the driver of the soil spectroscopy community to accelerate the pace of scientific discovery and innovation.

## Supporting information

**S1 Table.** Goodness-of-fit metrics from 10-fold cross-validation with refitting, calculated across different soil properties and model types: Root mean squared error (RMSE), mean error (bias), coefficient of determination (R^2^), Lin’s concordance correlation coefficient (CCC), ratio of performance to the interquartile range (RPIQ). Note: Except for `clay.tot_usda.a334_w.pctsilt.tot_usda.c62_w.pct`, `sand.tot_usda.c60_w.pct`, `ph.h2o_usda.a268_index`, and `ph.cacl2_usda.a481_index`, all the other soil properties’ metrics are reported in the natural logarithm space (with offset = 1, `log1p()` R function).

| Soil property | Model | RMSE | bias | $R^2$ | CCC | RPIQ |
| --- | --- | --- | --- | --- | --- | --- |
| acidity_usda.a795_cmolc.kg | mir_cubist_kssl_na.v1.2 | 0.26 | -0.01 | 0.93 | 0.96 | 4.38 |
| acidity_usda.a795_cmolc.kg | mir_cubist_oss1_na.v1.2 | 0.27 | -0.01 | 0.93 | 0.96 | 4.36 |
| acidity_usda.a795_cmolc.kg | nir.neospectra_cubist_oss1_na.v1.2 | 0.51 | -0.01 | 0.65 | 0.78 | 2.22 |
| acidity_usda.a795_cmolc.kg | visnir_cubist_oss1_na.v1.2 | 0.42 | 0.02 | 0.65 | 0.77 | 2.59 |
| aggstb_usda.a1_w.pct | mir_cubist_kssl_na.v1.2 | 0.64 | -0.04 | 0.69 | 0.80 | 2.92 |
| aggstb_usda.a1_w.pct | mir_cubist_oss1_na.v1.2 | 0.64 | -0.04 | 0.69 | 0.80 | 2.91 |

Continuation of S1 Table
| Soil property | Model | RMSE | bias | R<sup>2</sup> | CCC | RPIQ |
| --- | --- | --- | --- | --- | --- | --- |
| al.dith_usda.a65_w.pct | mir_cubist_kssl_na_v1.2 | 0.06 | 0.00 | 0.91 | 0.95 | 2.70 |
| al.dith_usda.a65_w.pct | mir_cubist_ossl_na_v1.2 | 0.06 | 0.00 | 0.91 | 0.95 | 2.72 |
| al.dith_usda.a65_w.pct | nir.neospectra_cubist_ossl_na_v1.2 | 0.09 | 0.00 | 0.74 | 0.84 | 1.49 |
| al.ext_usda.a1056_mg.kg | mir_cubist_kssl_na_v1.2 | 0.52 | -0.01 | 0.92 | 0.96 | 2.90 |
| al.ext_usda.a1056_mg.kg | mir_cubist_ossl_na_v1.2 | 0.38 | -0.02 | 0.92 | 0.96 | 2.69 |
| al.ext_usda.a69_cmolc.kg | mir_cubist_kssl_na_v1.2 | 0.23 | -0.00 | 0.90 | 0.95 | 4.80 |
| al.ext_usda.a69_cmolc.kg | mir_cubist_ossl_na_v1.2 | 0.23 | -0.00 | 0.90 | 0.94 | 4.74 |
| al.ext_usda.a69_cmolc.kg | nir.neospectra_cubist_ossl_na_v1.2 | 0.45 | 0.03 | 0.54 | 0.70 | 2.16 |
| al.ext_usda.a69_cmolc.kg | visnir_cubist_ossl_na_v1.2 | 0.43 | 0.02 | 0.63 | 0.76 | 2.66 |
| al.ox_usda.a59_w.pct | mir_cubist_kssl_na_v1.2 | 0.06 | 0.00 | 0.94 | 0.97 | 2.60 |
| al.ox_usda.a59_w.pct | mir_cubist_ossl_na_v1.2 | 0.06 | 0.00 | 0.94 | 0.97 | 2.60 |
| al.ox_usda.a59_w.pct | nir.neospectra_cubist_ossl_na_v1.2 | 0.10 | 0.01 | 0.77 | 0.85 | 1.17 |
| awc.33.1500kPa_usda.c80_w.frac | mir_cubist_kssl_na_v1.2 | 0.04 | 0.00 | 0.48 | 0.61 | 1.57 |
| awc.33.1500kPa_usda.c80_w.frac | mir_cubist_ossl_na_v1.2 | 0.04 | 0.00 | 0.48 | 0.61 | 1.58 |
| awc.33.1500kPa_usda.c80_w.frac | nir.neospectra_cubist_ossl_na_v1.2 | 0.05 | 0.00 | 0.17 | 0.31 | 1.37 |
| b.ext_mel3_mg.kg | mir_cubist_ossl_na_v1.2 | 0.17 | 0.01 | 0.80 | 0.88 | 1.65 |
| bd_usda.a4_g.cm3 | mir_cubist_kssl_na_v1.2 | 0.12 | -0.01 | 0.75 | 0.86 | 2.03 |
| bd_usda.a4_g.cm3 | mir_cubist_ossl_na_v1.2 | 0.12 | -0.01 | 0.75 | 0.86 | 2.04 |
| bd_usda.a4_g.cm3 | nir.neospectra_cubist_ossl_na_v1.2 | 0.08 | -0.01 | 0.45 | 0.62 | 1.46 |
| bd_usda.a4_g.cm3 | visnir_cubist_kssl_na_v1.2 | 0.14 | -0.01 | 0.66 | 0.79 | 1.78 |
| bd_usda.a4_g.cm3 | visnir_cubist_ossl_na_v1.2 | 0.13 | -0.01 | 0.67 | 0.79 | 1.79 |
| c.tot_usda.a622_w.pct | mir_cubist_kssl_na_v1.2 | 0.10 | 0.00 | 0.99 | 1.00 | 12.25 |
| c.tot_usda.a622_w.pct | mir_cubist_ossl_na_v1.2 | 0.10 | 0.00 | 0.99 | 1.00 | 11.76 |
| c.tot_usda.a622_w.pct | nir.neospectra_cubist_ossl_na_v1.2 | 0.27 | -0.01 | 0.77 | 0.87 | 2.89 |
| c.tot_usda.a622_w.pct | visnir_cubist_kssl_na_v1.2 | 0.24 | -0.01 | 0.97 | 0.98 | 10.54 |
| c.tot_usda.a622_w.pct | visnir_cubist_ossl_na_v1.2 | 0.23 | -0.01 | 0.97 | 0.98 | 10.91 |
| ca.ext_usda.a1059_mg.kg | mir_cubist_kssl_na_v1.2 | 0.43 | -0.02 | 0.94 | 0.97 | 4.78 |
| ca.ext_usda.a1059_mg.kg | mir_cubist_ossl_na_v1.2 | 0.41 | -0.02 | 0.94 | 0.97 | 5.36 |
| ca.ext_usda.a722_cmolc.kg | mir_cubist_kssl_na_v1.2 | 0.27 | -0.01 | 0.96 | 0.98 | 7.03 |

Continuation of S1 Table
| Soil property | Model | RMSE | bias | R<sup>2</sup> | CCC | RPIQ |
| --- | --- | --- | --- | --- | --- | --- |
| ca.ext_usda.a722_cmolc.kg | mir_cubist_ossl_na_v1.2 | 0.28 | -0.01 | 0.95 | 0.98 | 7.01 |
| ca.ext_usda.a722_cmolc.kg | nir.neospectra_cubist_ossl_na_v1.2 | 0.58 | -0.02 | 0.80 | 0.89 | 3.30 |
| ca.ext_usda.a722_cmolc.kg | visnir_cubist_ossl_na_v1.2 | 0.51 | -0.01 | 0.85 | 0.92 | 4.85 |
| caco3_usda.a54_w.pct | mir_cubist_kssl_na_v1.2 | 0.18 | -0.00 | 0.98 | 0.99 | 12.58 |
| caco3_usda.a54_w.pct | mir_cubist_ossl_na_v1.2 | 0.20 | -0.00 | 0.97 | 0.99 | 11.23 |
| caco3_usda.a54_w.pct | nir.neospectra_cubist_ossl_na_v1.2 | 0.65 | -0.02 | 0.67 | 0.80 | 3.06 |
| caco3_usda.a54_w.pct | visnir_cubist_kssl_na_v1.2 | 0.39 | 0.02 | 0.87 | 0.93 | 2.91 |
| caco3_usda.a54_w.pct | visnir_cubist_ossl_na_v1.2 | 0.36 | 0.01 | 0.91 | 0.95 | 3.14 |
| cec_usda.a723_cmolc.kg | mir_cubist_kssl_na_v1.2 | 0.16 | -0.00 | 0.97 | 0.98 | 6.68 |
| cec_usda.a723_cmolc.kg | mir_cubist_ossl_na_v1.2 | 0.17 | -0.00 | 0.96 | 0.98 | 6.12 |
| cec_usda.a723_cmolc.kg | nir.neospectra_cubist_ossl_na_v1.2 | 0.38 | -0.03 | 0.72 | 0.83 | 2.52 |
| cec_usda.a723_cmolc.kg | visnir_cubist_ossl_na_v1.2 | 0.34 | -0.02 | 0.80 | 0.88 | 2.94 |
| cf_usda.c236_w.pct | mir_cubist_kssl_na_v1.2 | 0.89 | 0.05 | 0.59 | 0.72 | 2.33 |
| cf_usda.c236_w.pct | mir_cubist_ossl_na_v1.2 | 0.90 | 0.05 | 0.58 | 0.71 | 2.31 |
| cf_usda.c236_w.pct | nir.neospectra_cubist_ossl_na_v1.2 | 1.16 | 0.11 | 0.33 | 0.48 | 2.29 |
| cf_usda.c236_w.pct | visnir_cubist_ossl_na_v1.2 | 0.67 | -0.04 | 0.33 | 0.47 | 1.65 |
| clay.tot_usda.a334_w.pct | mir_cubist_kssl_na_v1.2 | 3.95 | 0.02 | 0.94 | 0.97 | 5.69 |
| clay.tot_usda.a334_w.pct | mir_cubist_ossl_na_v1.2 | 5.64 | 0.10 | 0.90 | 0.94 | 4.20 |
| clay.tot_usda.a334_w.pct | nir.neospectra_cubist_ossl_na_v1.2 | 7.30 | 0.12 | 0.72 | 0.83 | 2.60 |
| clay.tot_usda.a334_w.pct | visnir_cubist_ossl_na_v1.2 | 6.61 | 0.27 | 0.82 | 0.89 | 3.02 |
| cu.ext_usda.a1063_mg.kg | mir_cubist_kssl_na_v1.2 | 0.25 | 0.00 | 0.86 | 0.92 | 4.25 |
| cu.ext_usda.a1063_mg.kg | mir_cubist_ossl_na_v1.2 | 0.29 | 0.01 | 0.78 | 0.87 | 3.23 |
| ec_usda.a364_ds.m | mir_cubist_kssl_na_v1.2 | 0.32 | 0.03 | 0.87 | 0.92 | 1.72 |
| ec_usda.a364_ds.m | mir_cubist_ossl_na_v1.2 | 0.31 | 0.03 | 0.86 | 0.92 | 1.69 |
| ec_usda.a364_ds.m | nir.neospectra_cubist_ossl_na_v1.2 | 0.36 | 0.06 | 0.55 | 0.65 | 0.70 |
| ec_usda.a364_ds.m | visnir_cubist_ossl_na_v1.2 | 0.12 | 0.02 | 0.62 | 0.73 | 1.28 |
| fe.dith_usda.a66_w.pct | mir_cubist_kssl_na_v1.2 | 0.15 | 0.00 | 0.91 | 0.95 | 4.40 |
| fe.dith_usda.a66_w.pct | mir_cubist_ossl_na_v1.2 | 0.15 | 0.00 | 0.91 | 0.95 | 4.48 |
| fe.dith_usda.a66_w.pct | nir.neospectra_cubist_ossl_na_v1.2 | 0.25 | -0.00 | 0.62 | 0.75 | 2.15 |

Continuation of S1 Table
| Soil property | Model | RMSE | bias | R<sup>2</sup> | CCC | RPIQ |
| --- | --- | --- | --- | --- | --- | --- |
| fe.ext.usda.a1064_mg.kg | mir_cubist_kssl_na_v1.2 | 0.42 | 0.01 | 0.80 | 0.89 | 2.60 |
| fe.ext.usda.a1064_mg.kg | mir_cubist_ossl_na_v1.2 | 0.39 | 0.00 | 0.76 | 0.85 | 2.38 |
| fe.ox.usda.a60_w.pct | mir_cubist_kssl_na_v1.2 | 0.12 | 0.01 | 0.82 | 0.89 | 2.75 |
| fe.ox.usda.a60_w.pct | mir_cubist_ossl_na_v1.2 | 0.13 | 0.01 | 0.80 | 0.88 | 2.65 |
| fe.ox.usda.a60_w.pct | nir.neospectra_cubist_ossl_na_v1.2 | 0.15 | 0.01 | 0.58 | 0.70 | 1.75 |
| k.ext.usda.a1065_mg.kg | mir_cubist_kssl_na_v1.2 | 0.52 | -0.01 | 0.81 | 0.88 | 2.14 |
| k.ext.usda.a1065_mg.kg | mir_cubist_ossl_na_v1.2 | 0.52 | -0.01 | 0.76 | 0.85 | 2.53 |
| k.ext.usda.a725_cmolc.kg | mir_cubist_kssl_na_v1.2 | 0.17 | 0.01 | 0.80 | 0.88 | 2.40 |
| k.ext.usda.a725_cmolc.kg | mir_cubist_ossl_na_v1.2 | 0.17 | 0.01 | 0.79 | 0.87 | 2.36 |
| k.ext.usda.a725_cmolc.kg | nir.neospectra_cubist_ossl_na_v1.2 | 0.22 | 0.03 | 0.54 | 0.68 | 1.75 |
| k.ext.usda.a725_cmolc.kg | visnir_cubist_ossl_na_v1.2 | 0.23 | 0.02 | 0.64 | 0.76 | 2.16 |
| mg.ext.usda.a1066_mg.kg | mir_cubist_kssl_na_v1.2 | 0.47 | -0.02 | 0.90 | 0.94 | 4.45 |
| mg.ext.usda.a1066_mg.kg | mir_cubist_ossl_na_v1.2 | 0.44 | -0.03 | 0.89 | 0.94 | 4.39 |
| mg.ext.usda.a724_cmolc.kg | mir_cubist_kssl_na_v1.2 | 0.26 | -0.00 | 0.92 | 0.96 | 5.28 |
| mg.ext.usda.a724_cmolc.kg | mir_cubist_ossl_na_v1.2 | 0.26 | -0.01 | 0.92 | 0.96 | 5.29 |
| mg.ext.usda.a724_cmolc.kg | nir.neospectra_cubist_ossl_na_v1.2 | 0.41 | 0.01 | 0.72 | 0.83 | 2.85 |
| mg.ext.usda.a724_cmolc.kg | visnir_cubist_ossl_na_v1.2 | 0.40 | 0.01 | 0.78 | 0.87 | 3.27 |
| mn.ext.usda.a1067_mg.kg | mir_cubist_kssl_na_v1.2 | 0.61 | -0.03 | 0.80 | 0.88 | 2.85 |
| mn.ext.usda.a1067_mg.kg | mir_cubist_ossl_na_v1.2 | 0.68 | -0.03 | 0.76 | 0.85 | 2.74 |
| mn.ext.usda.a70_mg.kg | mir_cubist_kssl_na_v1.2 | 0.74 | 0.05 | 0.69 | 0.79 | 2.06 |
| mn.ext.usda.a70_mg.kg | mir_cubist_ossl_na_v1.2 | 0.74 | 0.06 | 0.69 | 0.79 | 2.06 |
| mn.ext.usda.a70_mg.kg | nir.neospectra_cubist_ossl_na_v1.2 | 0.78 | 0.09 | 0.30 | 0.42 | 1.53 |
| n.tot.usda.a623_w.pct | mir_cubist_kssl_na_v1.2 | 0.06 | 0.00 | 0.97 | 0.98 | 3.82 |
| n.tot.usda.a623_w.pct | mir_cubist_ossl_na_v1.2 | 0.06 | 0.00 | 0.97 | 0.98 | 3.62 |
| n.tot.usda.a623_w.pct | nir.neospectra_cubist_ossl_na_v1.2 | 0.07 | 0.00 | 0.69 | 0.81 | 2.01 |
| n.tot.usda.a623_w.pct | visnir_cubist_kssl_na_v1.2 | 0.10 | -0.00 | 0.93 | 0.97 | 5.29 |
| n.tot.usda.a623_w.pct | visnir_cubist_ossl_na_v1.2 | 0.08 | 0.00 | 0.92 | 0.96 | 2.52 |
| na.ext.usda.a1068_mg.kg | mir_cubist_kssl_na_v1.2 | 0.78 | 0.03 | 0.75 | 0.85 | 2.64 |
| na.ext.usda.a1068_mg.kg | mir_cubist_ossl_na_v1.2 | 0.72 | 0.02 | 0.68 | 0.79 | 1.66 |

Continuation of S1 Table
| Soil property | Model | RMSE | bias | R<sup>2</sup> | CCC | RPIQ |
| --- | --- | --- | --- | --- | --- | --- |
| na.ext.usda.a726_cmolc.kg | mir_cubist_kssl_na_v1.2 | 0.40 | 0.03 | 0.82 | 0.89 | 0.48 |
| na.ext.usda.a726_cmolc.kg | mir_cubist_ossl_na_v1.2 | 0.42 | 0.04 | 0.80 | 0.87 | 0.47 |
| na.ext.usda.a726_cmolc.kg | nir.neospectra_cubist_ossl_na_v1.2 | 0.43 | 0.06 | 0.42 | 0.42 | 0.11 |
| na.ext.usda.a726_cmolc.kg | visnir_cubist_ossl_na_v1.2 | 0.34 | 0.04 | 0.54 | 0.65 | 0.78 |
| oc.usda.c729_w.pct | mir_cubist_kssl_na_v1.2 | 0.11 | 0.00 | 0.99 | 0.99 | 11.01 |
| oc.usda.c729_w.pct | mir_cubist_ossl_na_v1.2 | 0.12 | 0.00 | 0.99 | 0.99 | 9.98 |
| oc.usda.c729_w.pct | nir.neospectra_cubist_ossl_na_v1.2 | 0.25 | -0.01 | 0.81 | 0.89 | 3.22 |
| oc.usda.c729_w.pct | visnir_cubist_kssl_na_v1.2 | 0.23 | -0.01 | 0.97 | 0.99 | 11.24 |
| oc.usda.c729_w.pct | visnir_cubist_ossl_na_v1.2 | 0.24 | -0.00 | 0.94 | 0.97 | 4.20 |
| p.ext.usda.a1070_mg.kg | mir_cubist_kssl_na_v1.2 | 0.81 | 0.01 | 0.68 | 0.79 | 2.84 |
| p.ext.usda.a1070_mg.kg | mir_cubist_ossl_na_v1.2 | 0.80 | 0.01 | 0.67 | 0.79 | 2.79 |
| p.ext.usda.a1070_mg.kg | nir.neospectra_cubist_ossl_na_v1.2 | 1.16 | -0.01 | 0.30 | 0.45 | 1.80 |
| p.ext.usda.a270_mg.kg | mir_cubist_kssl_na_v1.2 | 0.76 | 0.03 | 0.73 | 0.83 | 3.02 |
| p.ext.usda.a270_mg.kg | mir_cubist_ossl_na_v1.2 | 0.76 | 0.04 | 0.72 | 0.82 | 2.97 |
| p.ext.usda.a274_mg.kg | mir_cubist_kssl_na_v1.2 | 0.70 | 0.02 | 0.64 | 0.76 | 2.59 |
| p.ext.usda.a274_mg.kg | mir_cubist_ossl_na_v1.2 | 0.74 | 0.02 | 0.61 | 0.74 | 2.49 |
| p.ext.usda.a274_mg.kg | visnir_cubist_ossl_na_v1.2 | 1.02 | -0.14 | 0.42 | 0.55 | 1.27 |
| ph.cacl2.usda.a481_index | mir_cubist_kssl_na_v1.2 | 0.36 | 0.01 | 0.93 | 0.96 | 6.74 |
| ph.cacl2.usda.a481_index | mir_cubist_ossl_na_v1.2 | 0.36 | 0.01 | 0.93 | 0.96 | 6.73 |
| ph.cacl2.usda.a481_index | nir.neospectra_cubist_ossl_na_v1.2 | 0.68 | 0.02 | 0.74 | 0.85 | 3.54 |
| ph.cacl2.usda.a481_index | visnir_cubist_ossl_na_v1.2 | 0.38 | 0.00 | 0.93 | 0.96 | 7.15 |
| ph.h2o.usda.a268_index | mir_cubist_kssl_na_v1.2 | 0.39 | 0.00 | 0.92 | 0.95 | 5.83 |
| ph.h2o.usda.a268_index | mir_cubist_ossl_na_v1.2 | 0.40 | 0.01 | 0.91 | 0.95 | 5.63 |
| ph.h2o.usda.a268_index | nir.neospectra_cubist_ossl_na_v1.2 | 0.64 | 0.01 | 0.75 | 0.85 | 3.39 |
| ph.h2o.usda.a268_index | visnir_cubist_ossl_na_v1.2 | 0.43 | -0.00 | 0.90 | 0.95 | 5.76 |
| s.tot.usda.a624_w.pct | mir_cubist_kssl_na_v1.2 | 0.07 | 0.01 | 0.93 | 0.96 | 0.64 |
| s.tot.usda.a624_w.pct | mir_cubist_ossl_na_v1.2 | 0.07 | 0.01 | 0.93 | 0.96 | 0.63 |
| s.tot.usda.a624_w.pct | nir.neospectra_cubist_ossl_na_v1.2 | 0.08 | 0.01 | 0.89 | 0.94 | 0.34 |
| s.tot.usda.a624_w.pct | visnir_cubist_kssl_na_v1.2 | 0.08 | 0.01 | 0.77 | 0.84 | 0.47 |

Continuation of S1 Table
| Soil property | Model | RMSE | bias | R<sup>2</sup> | CCC | RPIQ |
| --- | --- | --- | --- | --- | --- | --- |
| s.tot_usda.a624_w.pct | visnir_cubist_oss1_na_v1.2 | 0.08 | 0.01 | 0.75 | 0.83 | 0.45 |
| sand.tot_usda.c60_w.pct | mir_cubist_kssl_na_v1.2 | 7.48 | 0.01 | 0.93 | 0.96 | 6.53 |
| sand.tot_usda.c60_w.pct | mir_cubist_oss1_na_v1.2 | 8.33 | 0.00 | 0.92 | 0.96 | 5.84 |
| sand.tot_usda.c60_w.pct | nir.neospectra_cubist_oss1_na_v1.2 | 18.16 | 0.63 | 0.59 | 0.72 | 2.58 |
| sand.tot_usda.c60_w.pct | visnir_cubist_oss1_na_v1.2 | 15.12 | 0.04 | 0.68 | 0.80 | 2.84 |
| silt.tot_usda.c62_w.pct | mir_cubist_kssl_na_v1.2 | 6.63 | 0.05 | 0.90 | 0.94 | 4.60 |
| silt.tot_usda.c62_w.pct | mir_cubist_oss1_na_v1.2 | 7.31 | 0.05 | 0.88 | 0.93 | 4.40 |
| silt.tot_usda.c62_w.pct | nir.neospectra_cubist_oss1_na_v1.2 | 13.79 | 0.18 | 0.54 | 0.67 | 2.17 |
| silt.tot_usda.c62_w.pct | visnir_cubist_oss1_na_v1.2 | 11.25 | 0.16 | 0.67 | 0.79 | 2.49 |
| wr.1500kPa_usda.a417_w.pct | mir_cubist_kssl_na_v1.2 | 0.18 | -0.00 | 0.94 | 0.97 | 4.46 |
| wr.1500kPa_usda.a417_w.pct | mir_cubist_oss1_na_v1.2 | 0.19 | -0.00 | 0.94 | 0.97 | 4.41 |
| wr.1500kPa_usda.a417_w.pct | nir.neospectra_cubist_oss1_na_v1.2 | 0.36 | -0.02 | 0.65 | 0.78 | 2.13 |
| wr.1500kPa_usda.a417_w.pct | visnir_cubist_oss1_na_v1.2 | 0.43 | -0.04 | 0.61 | 0.73 | 1.85 |
| wr.33kPa_usda.a415_w.pct | mir_cubist_kssl_na_v1.2 | 0.22 | 0.00 | 0.83 | 0.91 | 2.26 |
| wr.33kPa_usda.a415_w.pct | mir_cubist_oss1_na_v1.2 | 0.22 | -0.00 | 0.83 | 0.90 | 2.27 |
| wr.33kPa_usda.a415_w.pct | nir.neospectra_cubist_oss1_na_v1.2 | 0.38 | -0.03 | 0.31 | 0.46 | 1.21 |
| wr.33kPa_usda.a415_w.pct | visnir_cubist_oss1_na_v1.2 | 0.35 | -0.04 | 0.57 | 0.70 | 1.66 |
| zn.ext_usda.a1073_mg.kg | mir_cubist_kssl_na_v1.2 | 0.36 | 0.02 | 0.74 | 0.83 | 2.47 |
| zn.ext_usda.a1073_mg.kg | mir_cubist_oss1_na_v1.2 | 0.35 | 0.03 | 0.60 | 0.72 | 1.73 |

**S2 Table.** Goodness-of-fit metrics of organic carbon (oc usda.c729 w.pct) from an independent evaluation of the mid-infrared (MIR) models calibrated with the Open Soil Spectral Library (OSSL) database. Statistics are reported for a dataset representing long-term research (LTR) trial sites. Metrics provided in original units after back-transformation.

| Source | Model type | Model subset | n | RMSE | bias | R<sup>2</sup> | CCC | RPIQ |
| --- | --- | --- | --- | --- | --- | --- | --- | --- |
| KSSL spectra | plsr | kssl | 162 | 0.38 | -0.22 | 0.95 | 0.92 | 2.05 |
| KSSL spectra | plsr | ossl | 162 | 0.36 | -0.21 | 0.95 | 0.93 | 2.18 |
| KSSL spectra | cubist | kssl | 162 | 0.27 | -0.16 | 0.95 | 0.95 | 2.91 |
| KSSL spectra | cubist | ossl | 162 | 0.22 | -0.10 | 0.96 | 0.97 | 3.53 |
| Woodwell spectra | plsr | kssl | 162 | 0.44 | -0.18 | 0.92 | 0.90 | 1.79 |
| Woodwell spectra | plsr | ossl | 162 | 0.45 | -0.22 | 0.92 | 0.90 | 1.74 |
| Woodwell spectra | cubist | kssl | 162 | 0.51 | -0.32 | 0.91 | 0.87 | 1.54 |
| Woodwell spectra | cubist | ossl | 162 | 0.24 | -0.06 | 0.94 | 0.96 | 3.24 |

## Acknowledgments

This study was funded by the USDA National Institute of Food and Agriculture Award # 2020-67021-32467. We are grateful to the USDA NRCS National Soil Survey Center — Kellogg Soil Survey Laboratory, ICRAF-World Agroforestry, ISRIC-World Soil Information, the Africa Soil Information Service (AfSIS) funded by the Bill and Melinda Gates Foundation, the European Soil Data Centre, and ETH Zurich, for publishing and providing high-quality data that the OSSL can build upon. The AI4SoilHealth project has received funding from the European Union’s Horizon Europe research and innovation program under grant agreement No. 101086179.

